# Cross-species activation of hydrogen cyanide production by a promiscuous quorum-sensing receptor promotes *Chromobacterium subtsugae* competition in a dual-species model

**DOI:** 10.1101/2022.11.01.514797

**Authors:** Cheyenne Loo, Pratik Koirala, Nathan C. Smith, Kara C. Evans, Saida Benomar, Isabelle R. Parisi, Anna Oller, Josephine R. Chandler

**Author notes:** Address correspondence to Josephine R. Chandler,. Contributed equally (Cheyenne Loo, Pratik Koirala).

## Abstract

Many saprophytic bacteria have LuxR-I-type acyl-homoserine lactone (AHL) quorum-sensing systems that may be important for competing with other bacteria in complex soil communities. LuxR AHL receptors specifically interact with cognate AHLs to cause changes in expression of target genes. Some LuxR-type AHL receptors have relaxed specificity and are responsive to non-cognate AHLs. These promiscuous receptors might be used to sense and respond to AHLs produced by other bacteria by eavesdropping. We are interested in understanding the role of eavesdropping during interspecies competition. The soil saprophyte *Chromobacterium subtsugae* has a single AHL circuit, CviR-I, which produces and responds to *N-*hexanoyl-HSL (C6-HSL). The AHL receptor CviR can respond to a variety of AHLs in addition to C6-HSL. In prior studies we have utilized a coculture model with *C. subtsugae* and another soil saprophyte *Burkholderia thailandensis*. Using this model, we previously showed that promiscuous activation of CviR by *B. thailandensis* AHLs provides a competitive advantage to *C. subtsugae*. Here, we show that *B. thailandensis* AHLs activate transcription of dozens of genes in *C. subtsugae*, including the *hcnABC* genes coding for production of hydrogen cyanide. We show that hydrogen cyanide production is population density-dependent and demonstrate that the cross-induction of hydrogen cyanide by *B. thailandensis* AHLs provides a competitive advantage to *C. subtsugae*. Our results provide new information on *C. subtsugae* quorum sensing and are the basis for future studies aimed at understanding the role of eavesdropping in interspecies competition.

**IMPACT STATEMENT:** In quorum sensing, population density-dependent changes in gene regulation are the result of a cytoplasmic transcription regulator binding to a quorum sensing signal. The signal-receptor interaction is considered to be specific to ensure fidelity of the system. However, some quorum-sensing receptor proteins have relaxed specificity and can recognize and respond to a range of signals. These promiscuous receptors might provide some benefit by enabling interspecies activation of quorum sensing by “eavesdropping,” although the potential benefits of eavesdropping are not well studied. The current study utilizes a dual-species laboratory competition model, where one species has a promiscuous signal receptor and can respond to signals produced by the other species. In our study, we identify the signals that enable quorum sensing cross talk and show that cross talk promotes competition by inducing hydrogen cyanide production. Our results highlight how quorum sensing-enabled interspecies cross talk might provide an advantage during competition and provide a new basis for understanding how receptor-signal pairs might evolve in natural environments.

**DATA SUMMARY:** The authors confirm all supporting data, code and protocols have been provided within the article or through supplementary data files.

## Introduction

Quorum sensing is a population density-dependent signaling system that relies on small diffusible signaling molecules. Once a critical population threshold is reached, the molecules interact with cytoplasmic transcription factors to cause activation of specific genes and coordinate gene expression across the population. In *Proteobacteria*, one type of quorum sensing is carried out by acyl-homoserine lactones (AHLs) signals. These molecules have a lactone ring and a fatty acid tail that varies in length from 4 to 20 carbons and can also have modifications at the third carbon position, which may be unsubstituted, hydroxylated, or have an oxo-substitution. Typical AHL quorum-sensing gene circuits consist of a LuxR-family signal receptor and a LuxI-family signal synthase, which produces a cognate signal. A bacterium may possess multiple LuxR-I circuits that typically produce and respond to different AHLs. Although the AHLs generally have similar structures, the receptors are thought to interact with AHLs in a highly specific and selective manner (1). However, there are exceptions such as the CviR receptor of *Chromobacterium subtsugae* (formerly *C. violaceum*), which can promiscuously detect and respond to a range of AHLs (2, 3). The existence of such receptors suggests receptor promiscuity might have certain advantages in some conditions.

Quorum sensing systems control different behaviors in different bacteria, such as the production of toxins, exoproducts, and biofilm matrix components. Many of these behaviors might provide a benefit to individuals within groups or complex communities. For example, toxin production might be important for competition with other strains or species of bacteria. There is an increasing interest in understanding the specific benefits of quorum sensing in dynamic polymicrobial communities. A major barrier to understanding quorum sensing in this context is the need for laboratory models that enable studies of mixed-strain and mixed-species interactions in a controlled setting. Such models are increasingly being utilized, and are providing significant new insights into the role and mechanisms of quorum sensing in mixed microbial communities (for a review, see (4)).

This study utilizes a laboratory dual-species model that was previously developed where two soil bacteria, *Burkholderia thailandensis* and *Chromobacterium subtsugae*, use AHL-dependent quorum sensing to compete with the other species (5) (Fig. 1). *B. thailandensis* produces a ribosome-targeting antibiotic called bactobolin (6). Bactobolin is regulated by a LuxR-I system called BtaR2-I2, which senses and responds to 3OHC8-HSL and 3OHC10-HSL (7). Both bactobolin and the BtaR2-I2 system are important for *B. thailandensis* to compete with *C. subtsugae* (5). In addition to BtaR2-I2, *B. thailandensis* has two other complete quorum-sensing circuits, BtaR1-I1 and BtaR3-I3, which produce and respond to C8-HSL and 3OHC10-HSL, respectively (8). *C. subtsugae* has a single LuxR-I-type system, CviR-I, which produces and responds to C6-HSL (2). CviR-I is important for competition with *B. thailandensis* (5). CviR is known to activate production of a purple pigmented antibiotic, violacein, however mutational disruption of violacein had no effect on competition (5). We also showed that CviR can be activated by *B. thailandensis* AHLs by “eavesdropping” to promote the competitive ability of *C. subtsugae* (5).

**Fig. 1.**
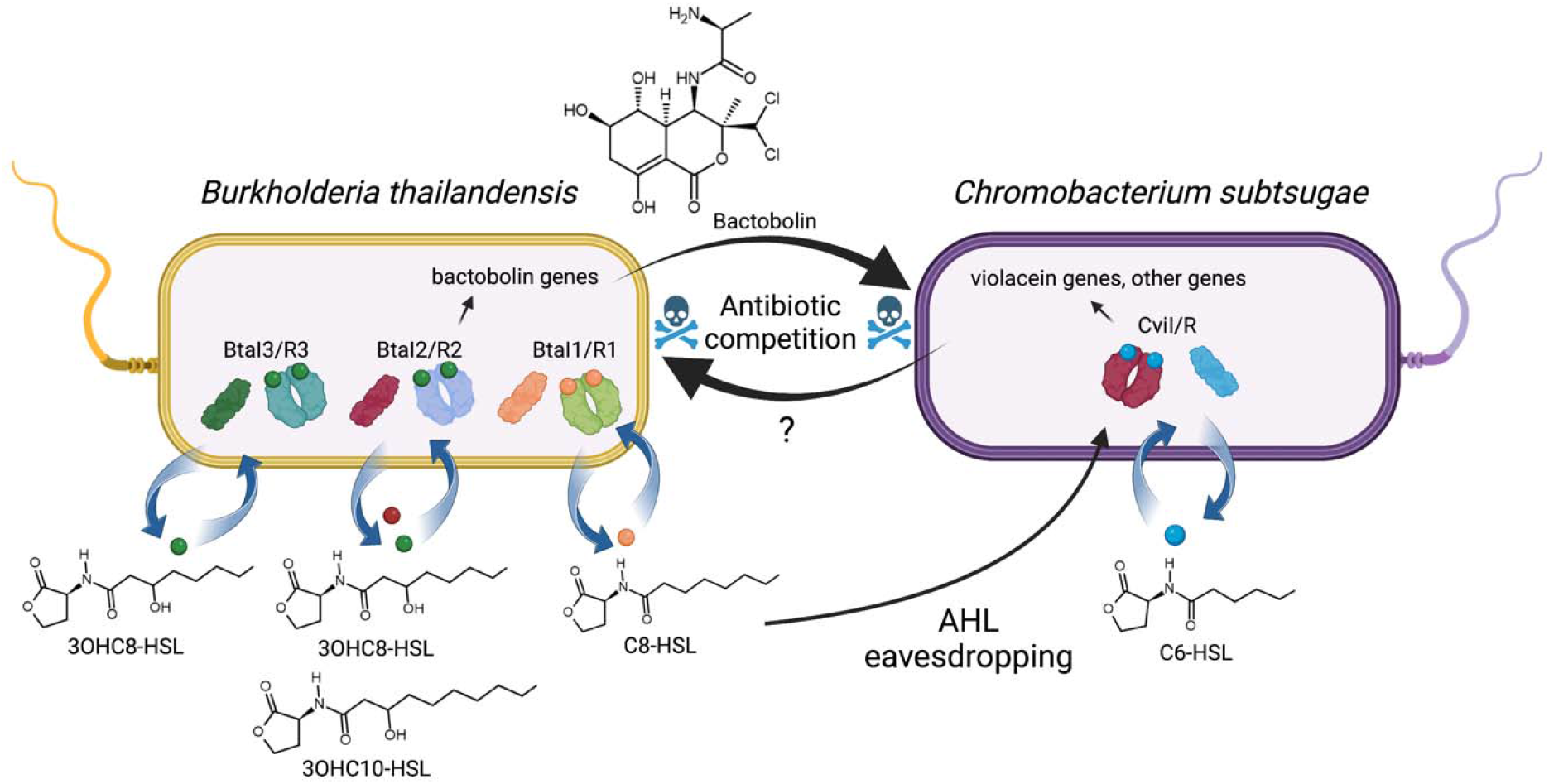
*Burkholderia thailandensis-Chromobacterium subtsugae* coculture model. The *B. thailandensis* quorum sensing system BtaR2-I2 produces and responds to 3OHC8-HSL and 3OHC10-HSL, and activates production of bactobolin antibiotic. *B. thailandensis* also has two other quorum sensing systems, BtaR1-I1 and BtaR3-I3, which produce C8-HSL and 3OHC8-HSL, respectively. BtaR2-I2 is important for *B. thailandensis* to compete with *C. subtsugae* due to activation of bactobolin prodution. The *C. subtsugae* quorum sensing system CviR-I produce and responds to C6-HSL. CviR-I is important for *C. subtsugae* to compete with *B. thailandensis*, through an unknown mechanism.

In this study, we identify C8-HSL as the *B. thailandensis* AHL best able to activate *C. subtsugae* production of quorum-dependent antimicrobials. We use RNAseq to identify the full set of *C. subtsugae* genes that can be activated by either C8-HSL or the native signal, C6-HSL. Among the genes most highly activated by both signals are those coding for hydrogen cyanide biosynthesis. We use our coculture model to validate the importance of hydrogen cyanide for *C. subtsugae* competition. We also demonstrate that the hydrogen cyanide genes can be transcriptionally activated by CviR and both native and non-native signals. The results provide new information on promiscuous AHL responses in *C. subtsugae* and further support the idea that AHLs drive important interactions through interspecies crosstalk.

## METHODS

### Bacterial culture conditions and reagents

All strains were grown in Lysogeny Broth (LB) broth (10g tryptone, 5g yeast extract, and 10g NaCl in 1L water), LB with morpholine-propanesulfonic acid (LB-MOPS, 50mM; pH 7), or in LB with 1.5% (wt/vol) Bacto-Agar. All broth cultures were incubated at 30 °C (*C. subtsugae* or *C. subtsugae-B. thailandensis* cocultures) or 37 °C (*B. thailandensis* or *E. coli*). Synthetic acyl-homoserine lactone signals were purchased from Cayman Chemicals (MI, USA) and stored in ethyl acetate acidified with 0.1% of glacial acetic acid. In all cases, synthetic signals were added to the culture flask and dried down completely using a stream of nitrogen gas prior to adding the culture media. For strain constructions, we used gentamicin at 50 μg ml^−1^ (*C. subtsugae*) or 15 μg ml^−1^ (*E. coli*). For selection from cocultures, we used gentamicin at 100 μg ml^−1^ (*B. thailandensis*) and trimethoprim at 100 μg ml^−1^ (*C. subtsugae*).

Genomic DNA, PCR and DNA fragments, and plasmid DNA were purified by using a Puregene Core A kit, plasmid purification miniprep kit, or PCR cleanup/gel extraction kits (Qiagen or IBI-MidSci) according to the manufacturer’s protocol. RNA was isolated using the Qiagen RNEasy Minikit.

### Bacterial strains and plasmids

Strains are summarized in Table 1. For cocultures, we used *B. thailandensis* strains JBT125 and BD20. Both are derived from strain E264 (9); BD20 is an bactobolin-deficient *btaK* mutant (7), and JBT125 is an AHL- and bactobolin-deficient *bta1-3, btaK* quadruple mutant (8). *C. subtsugae* strain CV017 (referred to as wild type, previously known as *C. violaceum* CV017) is a derivative of strain ATCC 31532 with a transposon insertion in gene CV_RS05185 causing overexpression of violacein (2). All *C. subtsugae* mutant strains were constructed from CV017 using allelic exchange and methods described below. For plasmid construction, we used *E. coli* strain DH10B (Invitrogen).

**Table 1.**
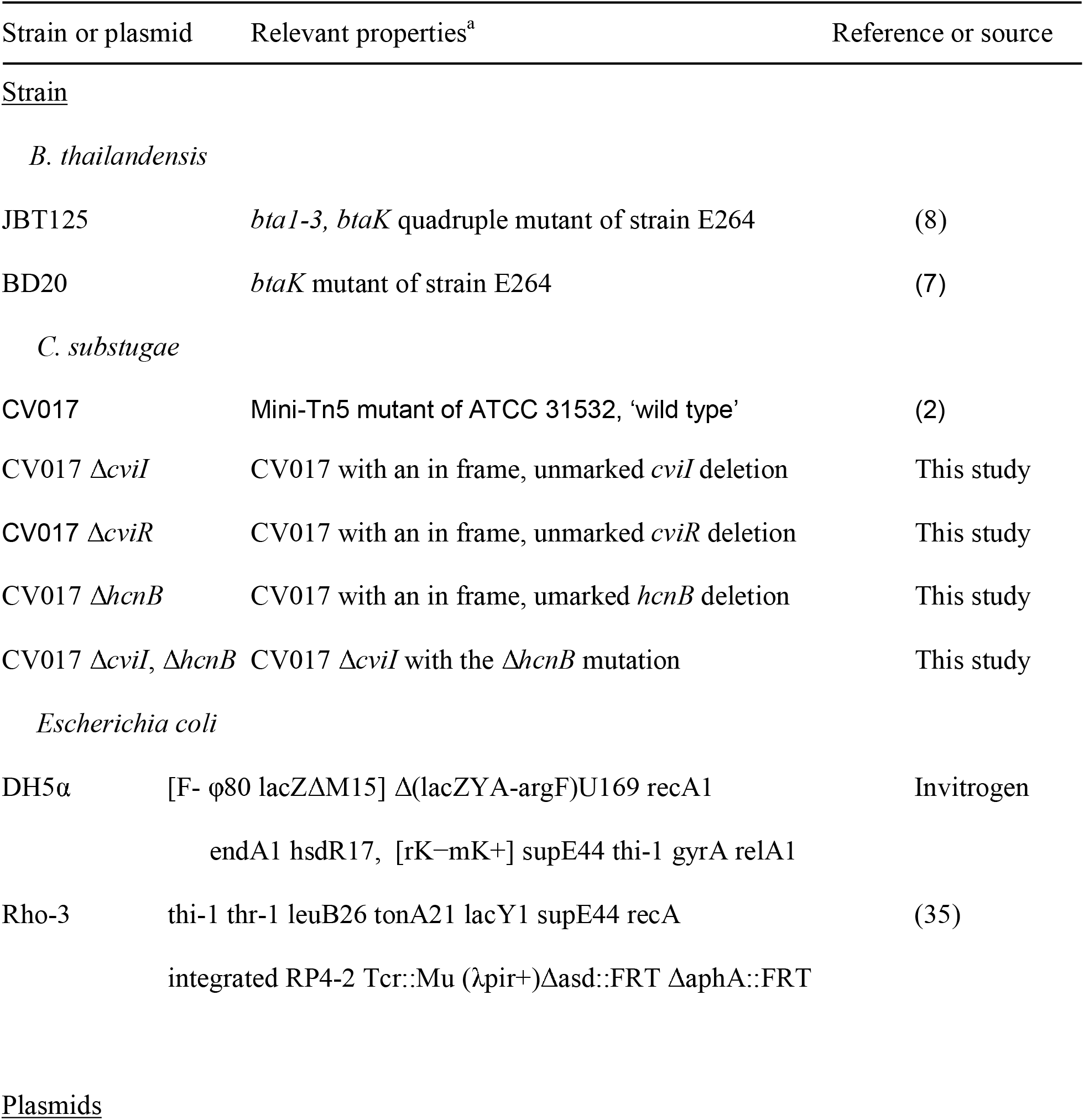
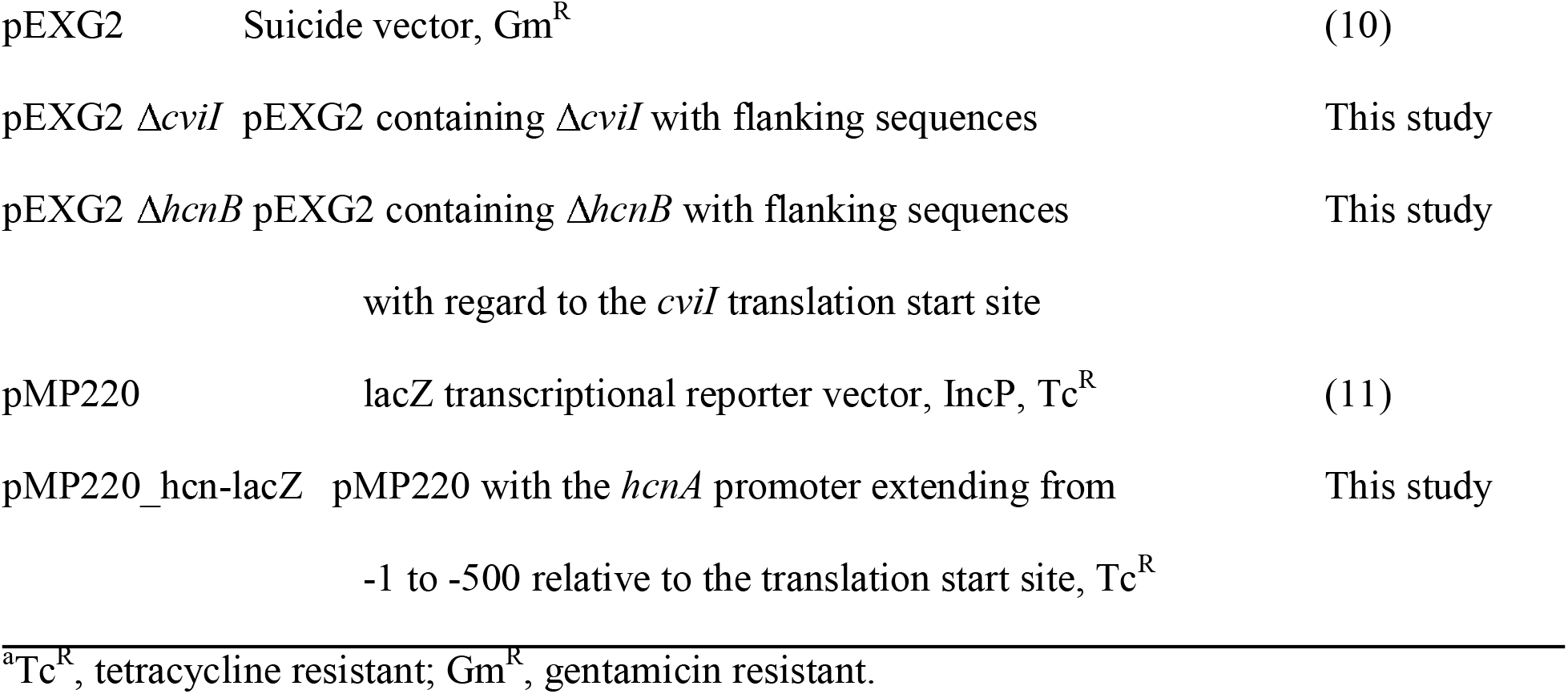
Bacterial strains and plasmids used in this study.

| Strain or plasmid | Relevant properties<sup>a</sup> | Reference or source |
| --- | --- | --- |
| <u>Strain</u> |  |  |
| *B. thailandensis* |  |  |
| JBT125 | *btaI-3*, *btaK* quadruple mutant of strain E264 | (8) |
| BD20 | *btaK* mutant of strain E264 | (7) |
| *C. substugae* |  |  |
| CV017 | Mini-Tn5 mutant of ATCC 31532, 'wild type' | (2) |
| CV017 $\Delta cviI$ | CV017 with an in frame, unmarked *cviI* deletion | This study |
| CV017 $\Delta cviR$ | CV017 with an in frame, unmarked *cviR* deletion | This study |
| CV017 $\Delta hcnB$ | CV017 with an in frame, unmarked *hcnB* deletion | This study |
| CV017 $\Delta cviI$, $\Delta hcnB$ | CV017 $\Delta cviI$ with the $\Delta hcnB$ mutation | This study |
| *Escherichia coli* |  |  |
| DH5α | [F- ϕ80 lacZΔM15] $\Delta$(lacZYA-argF)U169 recA1 | Invitrogen |
|  | endA1 hsdR17, [rK-mK+] supE44 thi-1 gyrA relA1 |  |
| Rho-3 | thi-1 thr-1 leuB26 tonA21 lacY1 supE44 recA | (35) |
| | integrated RP4-2 Tcr::Mu (λpir+) Δasd::FRT ΔaphA::FRT | |
| <u>Plasmids</u> |  |  |

|  |  |  |  |
| --- | --- | --- | --- |
| 450 | pEXG2 | Suicide vector, Gm<sup>R</sup> | (10) |
| 451 | pEXG2 Δ*cviI* | pEXG2 containing Δ*cviI* with flanking sequences | This study |
| 452 | pEXG2 Δ*hcnB* | pEXG2 containing Δ*hcnB* with flanking sequences | This study |
| 453 |  | with regard to the *cviI* translation start site |  |
| 454 | pMP220 | lacZ transcriptional reporter vector, IncP, Tc<sup>R</sup> | (11) |
| 455 | pMP220_*hcn*-lacZ | pMP220 with the *hcnA* promoter extending from | This study |
| 456 |  | -1 to -500 relative to the translation start site, Tc<sup>R</sup> |  |

<sup>a</sup>Tc<sup>R</sup>, tetracycline resistant; Gm<sup>R</sup>, gentamicin resistant.

Unmarked, in-frame deletions of *cviR, cviI*, and *hcnB* (Cv017_06925) were constructed using the following method. DNA fragments were generated by PCR or DNA synthesis (Genscript, New Jersey) containing ∼500-1000 bp DNA flanking each gene and fused together, creating an unmarked, non-polar deletion of each gene with incorporated XbaI and SacI restriction enzymes sites. These fragments were digested with XbaI and SacI and cloned into XbaI, SacI-digested pEXG2 (10). The relevant pEXG2 gene deletion plasmid was subsequently used to make the mutant *C. subtsugae* strain using methods described previously (5). Briefly, the relevant pEXG2 plasmid was transformed into *C. subtsugae* by electroporation or conjugation. Merodiploids were selected on LB agar containing gentamicin, and deletion mutants were counterselected on LB + 15% sucrose. Mutant strains were verified by testing for gentamicin sensitivity and by PCR-amplifying the deletion region and sequencing the PCR product.

For transcription reporter assays, plasmid pMP220 *hcnA-lacZ* was made by amplifying the *hcnA* promoter from *C. subtsugae* with primers hcn220R (5’-TATATCTAGAGATTAGCGGTTTGCCGTTCA-3’) and hcn220F (5’-TATATCTAGAGAAGCACCTTGCTGTGGTGAATG -3’), which incorporated the XbaI and KpnI sites. The amplified product was digested with XbaI and KpnI and ligated to XbaI- and KpnI-digested pMP220 (11). pMP220 *hcnA-lacZ* was introduced to *C. subtsugae* Δ*cviI* and Δ*cviR* by electroporation.

### RNA isolation

To isolate RNA for RNAseq, cultures were grown to an OD_600_ of 4.0 in 10 ml LB-MOPS in 125 ml non-baffled flasks. Bacteria (∼1 × 10^9^ cells) were removed from the cultures and suspended in RNAprotect bacteria reagent (Qiagen), pelleted by centrifugation, and stored at −80°C. Thawed cells were suspended in 1 ml RLT buffer (Qiagen) containing 2-mercaptoethanol and lysed by bead beating. RNA was purified using the RNAeasy minikit (Qiagen). Contaminating DNA was removed with Turbo DNase (Ambion), and RNA was obtained by using RNeasy MinElute cleanup kit (Qiagen).

### RNA-seq library construction, sequencing, mapping and analysis

Libraries for RNA-seq were prepared by Novogene using the Novogene mRNA-seq services project workflow. cDNA libraries were sequenced by Novogene using an Illumina platform and PE-150 reads. Sequences were aligned and mapped to the *C. subtsugae* CV017 genome (accession NZ_JAHDTB000000000) using Featurecounts and the Rsubread package in R (12). We determined differentially regulated genes for biological replicates using function edgeR (13) and using a false discovery rate [FDR] cutoff of 0.05. We proceeded with genes with >15 aligned reads and 2-fold or more regulation by C6- or C8-HSL relative to no AHL. Functions and their sources can be found at https://github.com/ChandlerLabKU/Cs_rnaseq. The data have been deposited in the NCBI sequence read archive (SRA database) (BioProject identification [ID] PRJNA894552). For Fig. 6, promoter sites and DNA recognition motifs were predicted using RSAT (14) and the transcription start site in the DNA region upstream of the *C. subtsugae hcnA* gene was predicted from the RNAseq data generated in this study using the TSSr program in R (15).

### Coculture experiments

Coculture experiments were conducted in 20 mL LB-MOPS medium in 125 mL non-baffled flasks. The inoculum was from logarithmic-phase pure cultures of *C. subtsugae* and *B. thailandensis*. The initial OD_600_ in the coculture was 0.05 for *B. thailandensis* (2—4x10^7^ cells per ml) and 0.005 for *C. subtsugae* (2 – 4x10^6^ cells per mL). After inoculating, cocultures were incubated at 30 ºC with shaking at 250 rpm for 24 hours. Colony forming units of each species were determined by using differential antibiotic selection on LB agar plates. *B. thailandensis* was selected with gentamicin and *C. subtsugae* was selected with trimethoprim.

### Transcription reporter assays

To assess activation of the *hcnA* promoter in *C. subtsugae*, overnight cultures of *C. subtsugae* strains carrying the pMP220_*hcn-lacZ* reporter plasmid were used as starters by diluting to an OD_600_ of 0.05. When experimental cultures reached an OD_600_ of 0.5 they were distributed to 2 ml Eppendorf tubes containing different concentrations of dried C6-HSL, C8-HSL, 3OHC8-HSL, and 3OHC10-HSL. The volume in each tube was 0.5 ml. After 5 h with shaking at 30°C, β-galactosidase activity was measured using the Tropix Galacto-Light Plus™ chemiluminescent kit according to the manufacturer’s protocol (Applied Biosystems, Foster City, CA).

#### Cyanide measurements

Cyanide concentrations were measured using <u>a CN^-^ ion-selective electrode (Cole-Parmer, USA). Cells were clarified from culture fluid using centrifugation followed by filter sterilization and the cell-free fluid was brought to a pH of 12 using NaOH prior to measugin the CN^-^ ion by conductivity. A standard curve using KCN was similarly measured and used to calculate the CN^-^ concentration in samples</u>.

## RESULTS

### *C. subtsugae* response to *B. thailandensis* AHLs in coculture

Previously, we showed that extracted fluid from *B. thailandensis* cultures containing all three *B. thailandensis* AHLs (C8-HSL, 3OHC8-HSL and 3OHC10-HSL) increase *C. subtsugae* competitiveness in a manner dependent on the *C. subtsugae* quorum-sensing receptor CviR (5). These results suggest that *C. subtsugae* CviR is responsive to one or several of the *B. thailandensis* AHLs. Therefore, we carried out coculture experiments to determine the individual contribution of each of the *B. thailandensis* AHL(s) to *C. subtsugae* competitiveness. We used signal synthase mutants of *C. subtsugae* and *B. thailandensis* for the competition experiment and added each of the *B. thailandensis* synthetic AHLs individually at 1 uM AHLs, which is a concentration in the range of that reported for similarly-grown pure cultures of *B. thailandensis* (16). The *B. thailandensis* strain JBT125 was also bactobolin-defective (Δ*btaK*) so that *C. subtsugae* would not be killed by AHL-induced bactobolin in the experiment. In our cocultures experiment we observed that the *C. subtsugae* native signal C6-HSL improved *C. subtsugae* competitiveness by ∼100-fold compared with the no signal condition (Fig. 2), similar to our prior study (5). Adding C8-HSL and 3OHC8-HSL also significantly improved competitiveness of *C. subtsugae* although to a lesser degree than the native C6-HSL. 3OHC10-HSL had a more variable effect than the other AHLs and was not significantly different from no AHLs, although this signal also appeared to improve competitiveness. Together, these results show that *C. subtsugae* is responsive to the *B. thailandensis* AHLs C8-HSL and 3OHC8-HSL, and possibly also to 3OHC10-HSL.

**Fig. 2.**
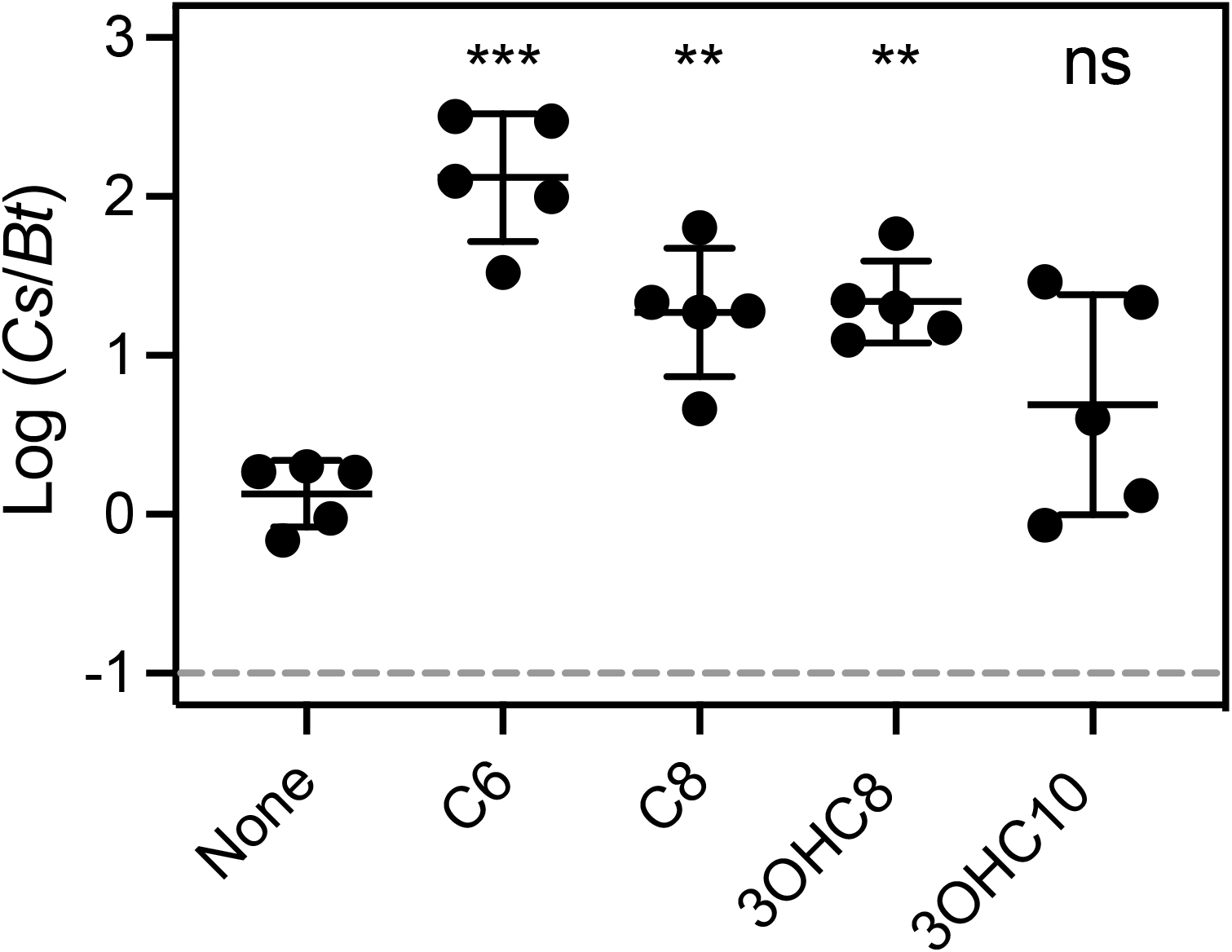
Contribution of *B. thailandensis* AHLs to *C. subtsugae* competitiveness. Cocultures were with *C. subtsugae* Δ*cviI* and *B. thailandensis* Δ*btaI1*, Δ*btaI2*, Δ*btaI3*, Δ*btaK* (JBT125). After 24 h of coculture growth, the ratio of *C. subtsugae* to *B. thailandensis* was determined by selective plating and colony counts. The dashed line indicates the initial ratio of *C. subtsugae* to *B. thailandensis*. The solid line represents the means and the vertical bars shows the standard error of the mean for each group. Synthetic AHLs were added at the beginning of the coculture to a final concentration of 1 µM. Statistical comparisons of each condition with no signal coculture were by one-way ANOVA; ***, *p*<0.001; **, *p*<0.005 and ns, not significant.

### Production of hydrogen cyanide is induced by quorum sensing and *hcnB*

We used RNAseq transcriptomic analysis to identify AHL-activated *C. subtsugae* genes as a first step to identify the factor important for competition with *B. thailandensis*. We compared transcripts in our Δ*cviI* strain grown with or without exogenously added synthetic AHLs. We grew cultures with either C6-HSL or C8-HSL, with the goal of identifying a set of genes with the ability to respond to both signals. We anticipated that the gene(s) required for *C. subtsugae* to compete via eavesdropping would be among this gene set. We collected cells for our analysis during the transition from logarithmic growth to stationary phase, which corresponds with an optical density at 600 nm (OD_600_) of 4.0. AHLs were added at 2 µM to ensure robust responses for detecting differences in gene expression.

Overall, 348 *C. subtsugae* genes were activated by C6-HSL, and 97 of these genes were also activated by C8-HSL (Table S1). In the set of genes most highly activated by both C6-HSL and C8-HSL (Table 2), we identified *hcnA, hcnB* and *hcnC*, which are predicted to code for biosynthesis of hydrogen cyanide. Hydrogen cyanide is important for *C. subtsugae* to protect itself from predation by other bacteria and kill mosquito larvae (17, 18), and is also important for *P. aeruginosa* to compete with *B. multivorans* in cocultures (19). Therefore, we focused our attention on hydrogen cyanide. To determine whether *C. subtsugae* produces hydrogen cyanide, we measured cyanide ion (CN^-^) in cell-free fluid from wild-type *C. subtsugae* cultures grown to various growth stages (early logarithmic, late logarithmic, stationary phase). The maximum concentration of CN^-^ was 3.3 ± 1.1 mM at an OD_600_ ∼8 (Fig. 3A). When adjusted for growth, the production of CN^-^ was population density-dependent (Fig. 3A), consistent with quorum sensing activation of these genes. To determine if either the *hcn* genes or quorum sensing was required for hydrogen cyanide production, we introduced a Δ*hcnB* mutation to the wild type genome and measured CN^-^ in culture fluid of the Δ*hcnB* and Δ*cviR* mutants and wild type. Growth-adjusted CN^-^ levels were low in both Δ*hcnB* and Δ*cviR* culture fluid compared with that of wild type (Fig. 3B). These results support the conclusion that *hcnB* and quorum sensing are important for hydrogen cyanide production in *C. subtsugae*.

**Table 2.**
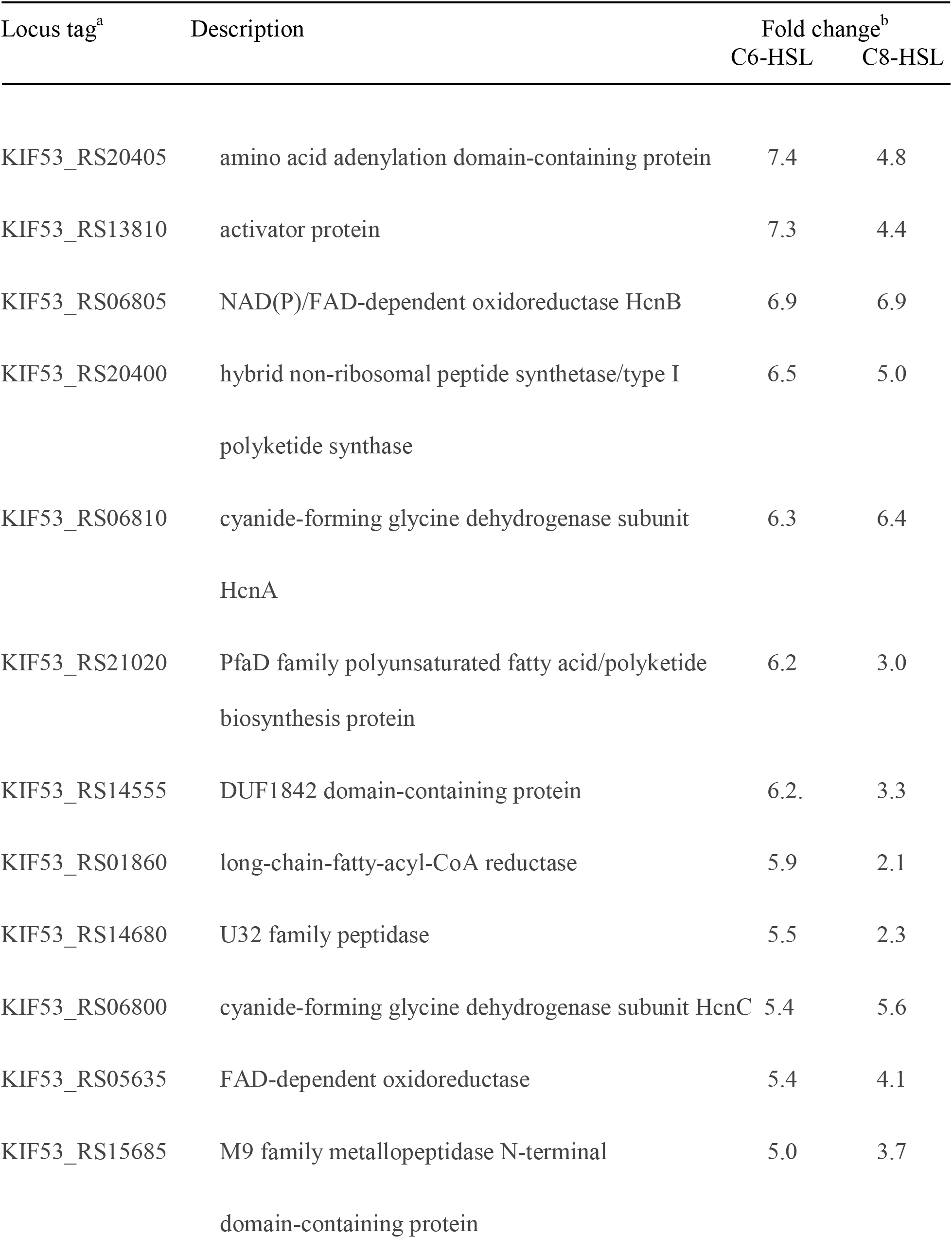
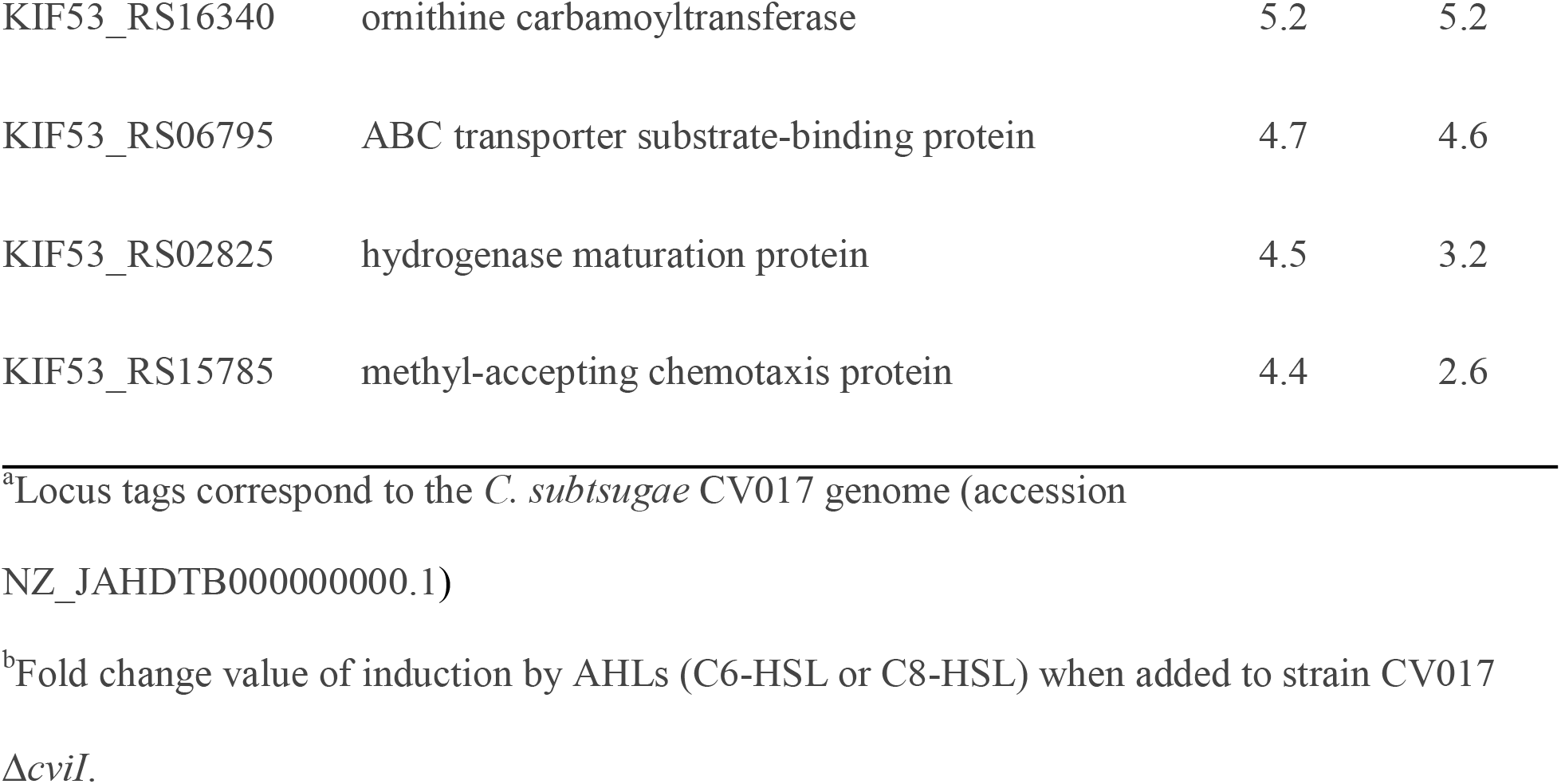
Genes most highly activated by C6-HSL and C8-HSL

| Locus tag<sup>a</sup> | Description | Fold change<sup>b</sup> |  |
| --- | --- | --- | --- |
|  |  | C6-HSL | C8-HSL |
| KIF53_RS20405 | amino acid adenylation domain-containing protein | 7.4 | 4.8 |
| KIF53_RS13810 | activator protein | 7.3 | 4.4 |
| KIF53_RS06805 | NAD(P)/FAD-dependent oxidoreductase HcnB | 6.9 | 6.9 |
| KIF53_RS20400 | hybrid non-ribosomal peptide synthetase/type I | 6.5 | 5.0 |
|  | polyketide synthase |  |  |
| KIF53_RS06810 | cyanide-forming glycine dehydrogenase subunit | 6.3 | 6.4 |
|  | HcnA |  |  |
| KIF53_RS21020 | PfaD family polyunsaturated fatty acid/polyketide | 6.2 | 3.0 |
|  | biosynthesis protein |  |  |
| KIF53_RS14555 | DUF1842 domain-containing protein | 6.2 | 3.3 |
| KIF53_RS01860 | long-chain-fatty-acyl-CoA reductase | 5.9 | 2.1 |
| KIF53_RS14680 | U32 family peptidase | 5.5 | 2.3 |
| KIF53_RS06800 | cyanide-forming glycine dehydrogenase subunit HcnC | 5.4 | 5.6 |
| KIF53_RS05635 | FAD-dependent oxidoreductase | 5.4 | 4.1 |
| KIF53_RS15685 | M9 family metallopeptidase N-terminal | 5.0 | 3.7 |
|  | domain-containing protein |  |  |

|  |  |  |  |  |
| --- | --- | --- | --- | --- |
| 482 | KIF53_RS16340 | ornithine carbamoyltransferase | 5.2 | 5.2 |
| 483 | KIF53_RS06795 | ABC transporter substrate-binding protein | 4.7 | 4.6 |
| 484 | KIF53_RS02825 | hydrogenase maturation protein | 4.5 | 3.2 |
| 485 | KIF53_RS15785 | methyl-accepting chemotaxis protein | 4.4 | 2.6 |

<sup>a</sup>Locus tags correspond to the *C. subtsugae* CV017 genome (accession
NZ\_JAHDTB000000000.1)
<sup>b</sup>Fold change value of induction by AHLs (C6-HSL or C8-HSL) when added to strain CV017
*ΔcviI*.

**Fig. 3.**
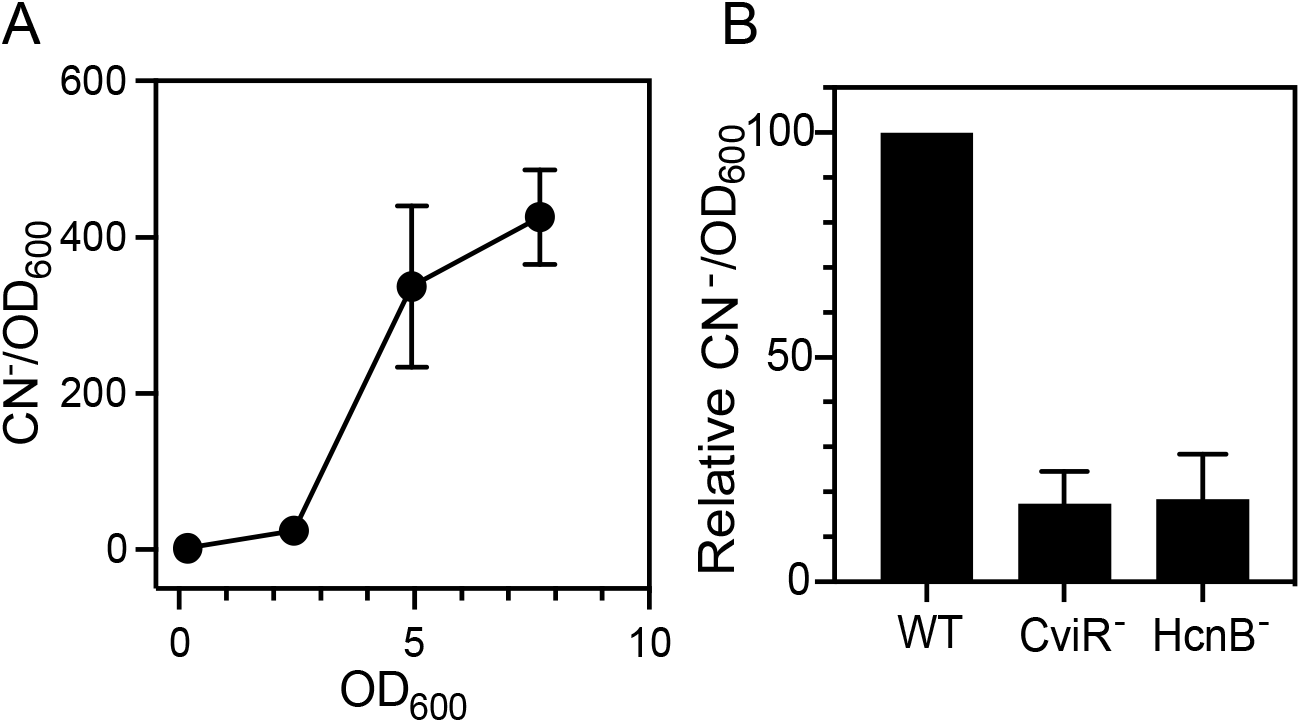
Cyanide measurements in *C. subtsugae* culture fluid. Cyanide (CN^-^) was measured in culture fluid of *C. subtsugae*. A. Wild type *C. subtsugae* (CV017) was grown to different population densities as indicated by the optical density at 600 nm (OD_600_) prior to measuring CN^-^. Data are shown as growth-adjusted CN^-^. The total CN^-^ concentration at OD_600_ of ∼8 corresponds with 3.3 ± 1.1 mM. B. The indicated *C. subtsugae* strain was grown for 12 h, which corresponds with OD_600_ of ∼8, prior to measuring population density and CN^-^. Data are shown as the growth-adjusted CN^-^ as a percentage of that of wild type. In all cases, data represent the average of 3 biological replicates and the vertical bars represents the standard error of the mean.

### Cyanide is important for *C. subtsugae* to compete with *B. thailandensis*

Because expression of the hydrogen cyanide biosynthesis genes is responsive to both C6-HSL and C8-HSL, we hypothesized that *B. thailandensis* AHLs increase competitiveness of *C. substugae* by activating hydrogen cyanide production through eavesdropping. As a first test of this hypothesis, we determined the susceptibility of *B. thailandensis* strain BD20 to a range of concentrations of potassium cyanide (0-750 uM), which mimics cyanide ions (20). We determined the 50% lethal dose (LD_50_) of potassium cyanide for *B. thailandensis* BD20 to be 119 µM (Fig. 4A). Thus, the concentration of cyanide produced by *C. subtsugae* (∼3 mM, Fig. 2A) is more than sufficient to kill *B. thailandensis*.

**Fig. 4.**
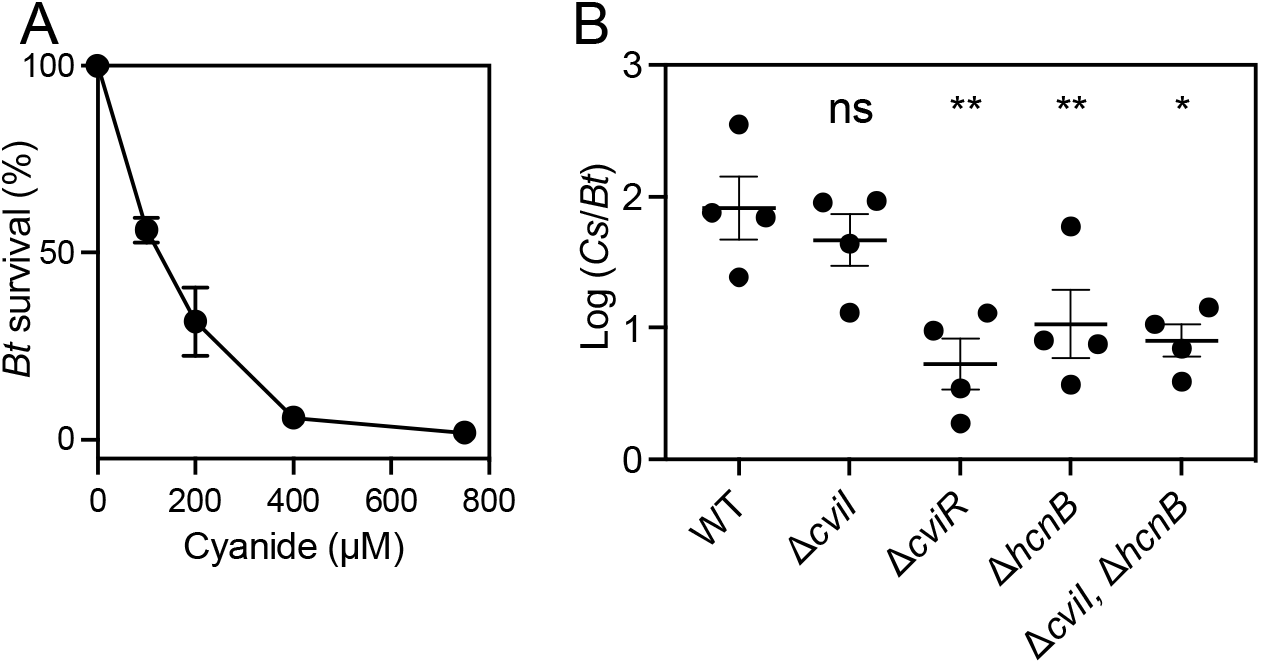
Cyanide toxicity and role in cocultures. A. Survival of *B. thailandensis* BD20 grown in the presence of different concentrations of cyanide for 8 hr from an initial OD_600_ of 0.005. The population density was measured and *B. thailandensis* (Bt) survival was determined as a percent of untreated culture grown under identical conditions. B. Cocultures were with *B. thailandensis* Δ*btaK* (BD20) and the *C. subtsugae* strain indicated. After 24 h of coculture growth, the ratio of *C. subtsugae* to *B. thailandensis* was determined by selective plating and colony counts. The initial ratio of *C. subtsugae* to *B. thailandensis* was 10^−1^. The solid line represents the means and the vertical bars shows the standard error for each group. Statistical comparisons of each condition compared with no signal were by one-way ANOVA; **, *p*<0.005; * = p < 0.05 and ns, not significant.

Next, we investigated the effects of hydrogen cyanide in cocultures (Fig. 4B). We competed the bactobolin-deficient *B. thailandensis* (BD20) with different genetic mutants of *C. subtsugae*. In these conditions, *B. thailandensis* BD20 produces AHLs that can activate the quorum-sensing system of *C. subtsugae* (5). Consistent with prior results (5), in these conditions the *C. subtsugae* Δ*cviI* mutant and wild type are similarly competitive, because the Δ*cviI* mutant has CviR that can be activated by *B. thailandensis* AHLs. The Δ*cviR* mutant cannot respond to any AHLs and, as expected, was significantly less competitive than the wild-type and Δ*cviI* mutant. To determine the role of hydrogen cyanide in competition, we tested the competitive ability of the Δ*hcnB* mutant. The Δ*hcnB* single mutant competed poorly with BD20 and was similar to the Δ*cviR* mutant. These results support that hydrogen cyanide is important for the competitive ability of *C. subtsugae*. We also tested the Δ*cviI*, Δ*hcnB* double mutant. In this strain, the competitive ability of *C. subtsugae* relies on AHLs produced by *B. thailandensis*. The Δ*cviI*, Δ*hcnB* double mutant also competed poorly, similar to the Δ*cviR* and Δ*hcnB* single mutants. These results support that hydrogen cyanide is important for *C. subtsugae* to compete in response to *B. thailandensis* AHLs. Together, the results support the idea that hydrogen cyanide is required for the competitive advantage provided to *C. subtsugae* in response to native AHLs and non-native AHLs.

### Sensitivity of *hcnA* promoter to AHLs

The hydrogen cyanide biosynthesis genes *hcnA, hcnB* and *hcnC* (the *hcn* genes) are adjacently encoded in the genome facing the same direction. There is a single putative promoter upstream of *hcnA*. We hypothesized that this promoter is activated either directly or indirectly by CviR in response to the *C. subtsugae-*produced signal C6-HSL, as well as the *B. thailandensis* signals C8-HSL, 3OHC8-HSL, and possibly 3OHC10-HSL. To test our hypothesis, we created a plasmid with a DNA fragment containing the putative promoter region upstream of *hcnA* fused to a promoterless *lacZ* gene. This plasmid was introduced into the *C. subtsugae* Δ*cviI* and Δ*cviR* mutants. The *hcnA-lacZ* reporter was activated by 5 µM of each of the four AHLs (C6-, C8-, 3OHC8- and 3OHC10-HSL) in the Δ*cviI* mutant, but not in the Δ*cviR* mutant (Fig. 5A). Thus, activation of *hcnA-lacZ* required CviR and AHLs. We also determined the signal sensitivity of the *hcnA-lacZ* reporter by testing each AHL at a range of concentrations (Fig. 5B). The *hcnA-lacZ* reporter responded most sensitively to the native signal C6-HSL. There was also a response to C8-HSL. The other two AHLs, 3OHC8-HSL and 3OHC10-HSL, activated the reporter only at the highest signal concentrations (1-5 µM). These results were consistent with the coculture results (Fig. 2), and demonstrate that C8-HSL and, to a lesser extent 3OHC8-HSL and 3OHC10-HSL, can activate the *hcn* gene promoter in a manner dependent on CviR.

**Fig. 5.**
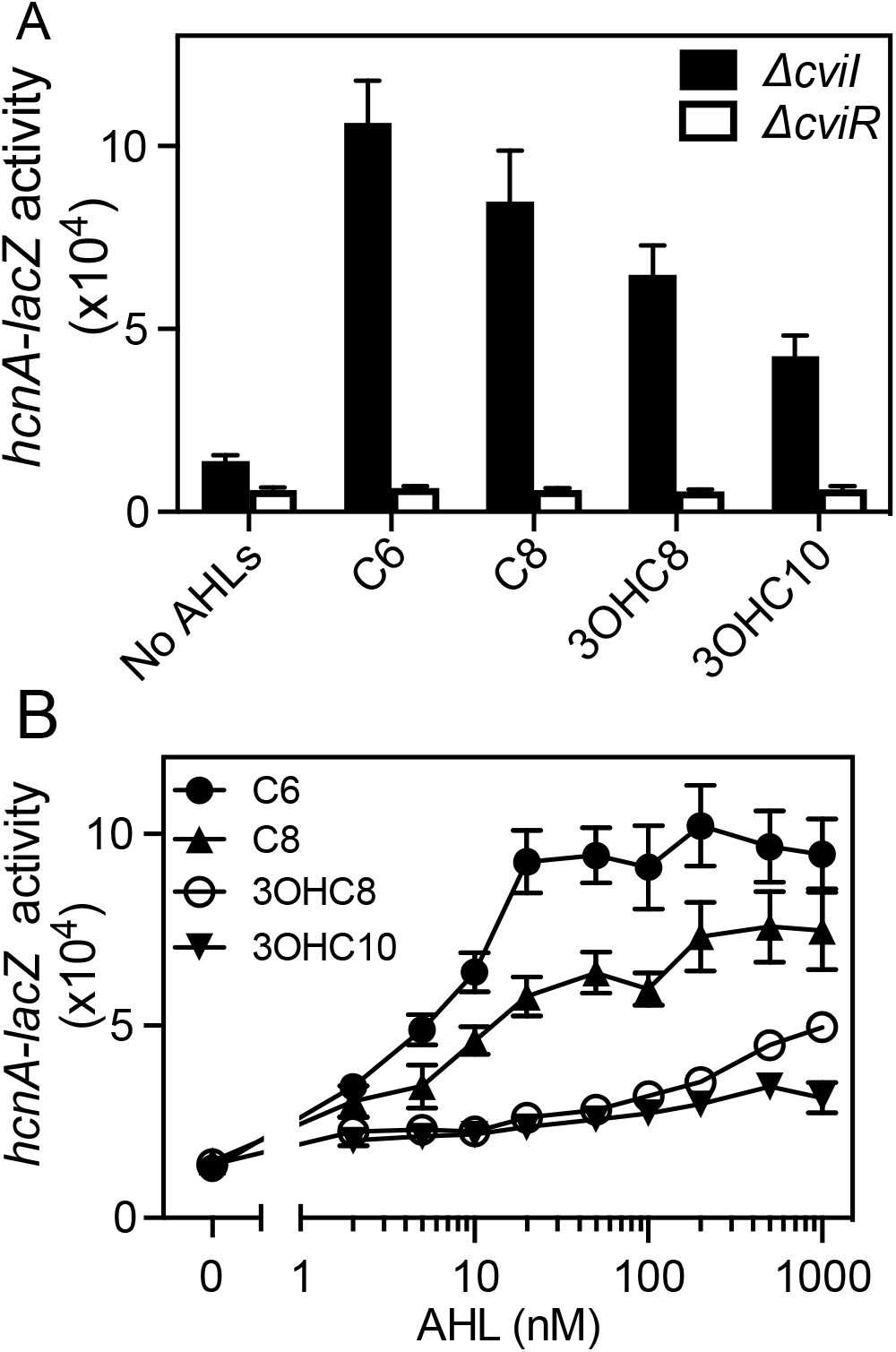
Sensitivity of the *hcnA* promoter to native and non-native AHLs. A. *C. subtsugae* Δ*cviI* or Δ*cviR* mutant harboring the *hcnA-lacZ* reporter was treated with 5 µM of AHL (C6-HSL, C8-HSL, 3OHC8-HSL or 3OHC10-HSL). B. *C. subtsugae* Δ*cviI* harboring the *hcnA-lacZ* reporter was treated with AHLs at the indicated concentrations. Results show the average of four independent experiments from two separate days, and the vertical bars represent the standard error.

**Fig. 6.**
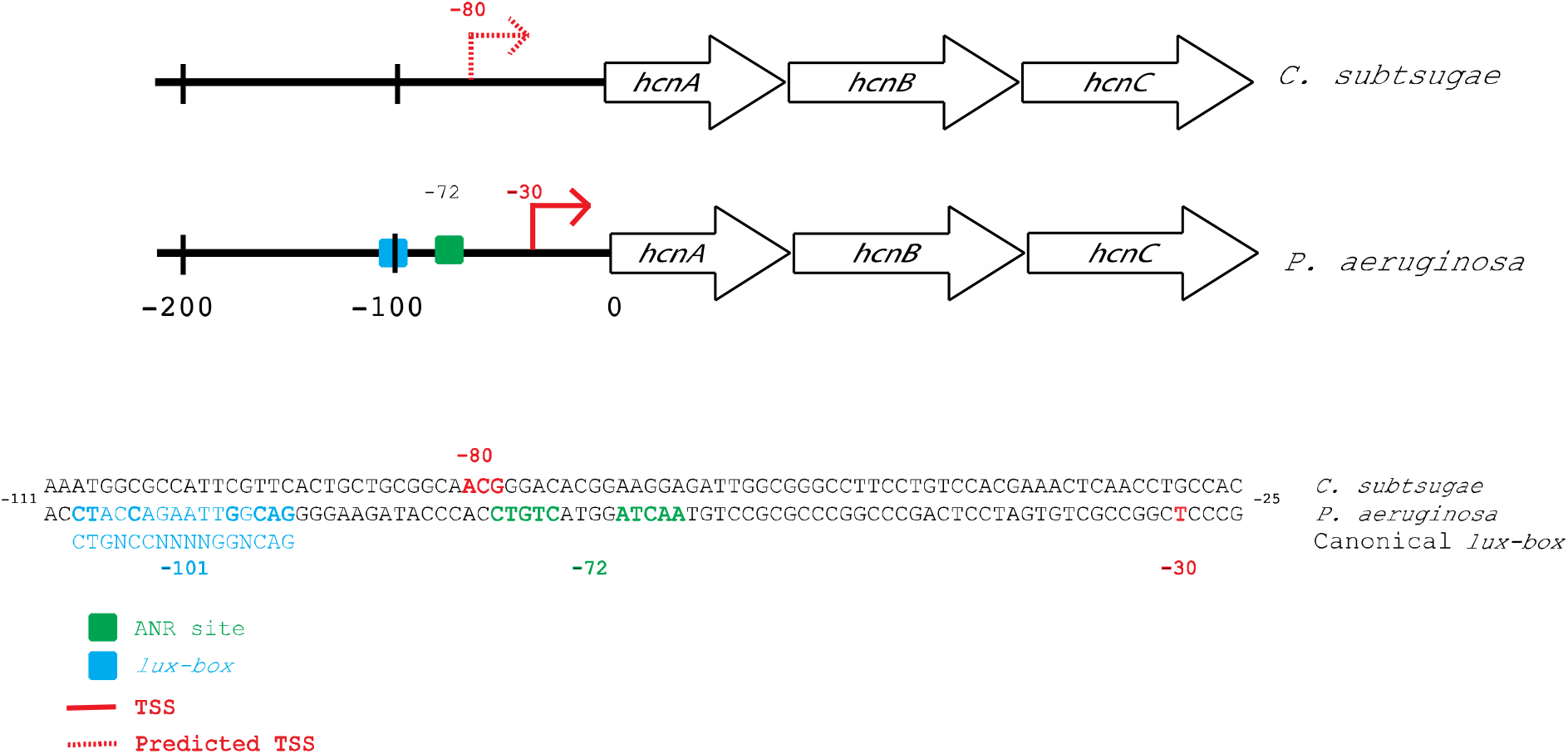
Hydrogen cyanide biosynthesis genes (*hcnABC*) in *C. subtsugae* and *P. aeruginosa*. Red arrows and lettering indicate the transcription start site. The arrow is dashed for *C. subtsugae* as it was predicted to be between nucleotides -79 to -81 (see Methods). Green boxes and lettering indicate the binding site for the anaerobic regulator (Anr) and the blue box and lettering indicate the *lux* box of *P. aeruginosa*, and the canonical lux box. Bolding in the *P*.*aeruginosa lux* box sequence indicate nucleotides that are conserved with the canonical *lux* box. No *lux* box or Fnr/Anr like sequences were identified in the *C. subtsugae* sequence (see Discussion text).

## DISCUSSION

The *B. thailandensis-C. subtsugae* model system was developed previously to examine the effects of quorum sensing and AHL-dependent eavesdropping in competition (5). In those studies, we elucidated the importance of *B. thailandensis* bactobolin for competition, however, the *C. subtsugae* toxin remained elusive. Here, we identified hydrogen cyanide as the *C. substugae* toxin, filling in this gap in knowledge. We have also shown that noncognate AHLs can induce dozens of genes in addition to those coding for hydrogen cyanide. Results with our coculture model support the idea that promiscuous receptors might enable interspecies interactions by coordinating broad changes in gene expression in response to AHLs from other species. Results of our studies also expand the list of genes known to be activated by the CviR-I system in *C. subtsugae*.

The *C. substugae* CviR receptor can activate certain genes in response to a wide range of AHLs (2). Although detailed structure-function studies of CviR and different AHLs have been carried out (21, 22), few studies have addressed the potential benefits of AHL receptor promiscuity for the cell. Receptor activation by non-self AHLs could enable detection of other bacteria and might provide certain benefits for interspecies competition (5) or cooperation (23). Indeed, there is evidence of eavesdropping in natural polymicrobial communities in plants (23-25), and eavesdropping has also been shown to occur in communities relevant to infections (26). Recent studies also support the idea that promiscuity might be a feature of a wider range of receptors than previously recognized (27). These studies support the idea that AHL-dependent cross talk might be an important but underrecognized facet of quorum sensing communication and that it warrants further study.

We find it interesting that in our study, the majority of genes most strongly activated by both C6-HSL and C8-HSL in *C. subtsugae* are predicted to code for antibiotic production (Table 2). The receptor BtaR2 in *B. thailandensis* is also highly promiscuous and regulates production of a potent and broad-spectrum antibiotic, bactobolin (6, 7). The role of these receptors in regulating antibiotic production supports the idea that promiscuous activation of receptors could play an important role in competition. The promiscuous activation of CviR by C8-HSL leads to the induction of a subset of genes that overlaps with that controlled by C6-HSL. It may be that these genes are more highly sensitive to CviR, enabling response to CviR activation by non-native signals. An alternative possibility is that C8-HSL changes CviR recognition of gene promoters, possibly through allosteric changes in the DNA binding motif. Future studies focused on the regulation of particular genes by different AHLs could help to elucidate the mechanism of CviR activation of gene promoters by non-native AHLs.

Prior studies show that receptor selectivity falls on a continuum from highly selective (e.g. *P. aeruginosa* RhlR) to intermediate (e.g. *P. aeruginosa* LasR) to highly promiscuous (e.g. CviR, and *B. thailandensis* BtaR2) (27). The affinity of receptors for a particular AHL can be influenced by changing specific residues within the AHL binding pocket of the receptor (21, 28-30). For example, although CviR is normally inhibited by the noncognate signal C10-HSL, replacing methionine at position 89 with serine converts this receptor to become C10-HSL-responsive (21). These results suggest that a particular receptor could easily adapt to become more or less promiscuous dependent on the benefits of promiscuity. In addition, receptor selectivity is influenced by receptor expression levels (27). Although very little is known about the environmental conditions that regulate quorum sensing receptors, one can imagine that conditions that increase receptor levels in the cell could increase AHL promiscuity. For example, it was recently demonstrated that the CviR receptor is activated by certain antibiotics (31), although the effects of antibiotics on receptor selectivity remain to be tested.

The results of this study are consistent with prior studies (17, 19, 32) supporting a role for hydrogen cyanide in competition. In *P. aeruginosa*, the hydrogen cyanide biosynthesis genes are part of an operon driven by a promoter upstream of *hcnA* (see Fig. 6). The *P. aeruginosa hcnA* promoter is quorum-sensing controlled, and has an experimentally validated binding site for the quorum sensing receptor LasR at position -101 relative to the translation start site (33). This region also has a binding site for the anaerobic regulator (Anr) at position -72 (33, 34) (Fig. 6). The intergenic region upstream of *hcnA* in *P. aeruginosa* and *C. substugae* share 45% identity; however, we were unable to find evidence of a *lux* box or Anr binding site anywhere in the *C. substugae hcnA* promoter. It is possible that CviR can recognize the *hcnA* promoter in the absence of a recognizable *lux* box, which has been reported in some cases for LasR (33). It is also possible that CviR controls the *hcn* genes indirectly. Either way, our results provide additional support for the idea that CviR can activate production of hydrogen cyanide in response to AHL from other species, supporting the idea that some quorum sensing-dependent competitive behaviors might be induced by interspecies cross talk.

## AUTHOR STATEMENTS

### Conflict of interest

The authors declare no conflicts of interest.

### Funding information

This work was supported by the National Institutes of Health (NIH) R35GM133572 to J.R.C. S.B. and C.L. were supported by NIH K-INBRE fellowships (P20GM103418), K.C.E. was supported by the NIH Chemical Biology Training Program (T32 GM08545), and P.K. was supported by the KU Weaver Fellowship. We also thank the KU Genomics Core, which is supported by NIH grants P20GM103638 (COBRE CMADP) and P20GM103418 (K-INBRE).

## Acknowledgements

We acknowledge Drs. Ajai Dandekar and Amy Schaefer for helpful discussions and Nicole Smalley for technical input. BioRender was used for the creation of Fig. 1.

## Notes

### Competing Interest Statement

The authors have declared no competing interest.

### Summary of Updates

This version of the manuscript has been revised in response to reviewer criticisms as part of the publication process.

