## Supplemental Information for "Cross-species activation of hydrogen cyanide production by a promiscuous quorum-sensing receptor promotes *Chromobacterium subtsugae* competition in a dual-species model"

### *C. subtsugae* genes activated by C6-HSL

| Locus tag corresponding with accession NZ_JAHDTB000000000.1 | Predicted gene function | $Log_2$ fold change compared with no AHLs |
|---|---|---|
| KIF53_RS20405 | amino acid adenylation domain-containing protein | 7.43 |
| KIF53_RS13810 | activator protein | 7.28 |
| KIF53_RS06805 | NAD(P)/FAD-dependent oxidoreductase | 6.86 |
| KIF53_RS20410 | SDR family NAD(P)-dependent oxidoreductase | 6.79 |
| KIF53_RS01865 | acyl-protein synthase | 6.76 |
| KIF53_RS20400 | polyketide synthase | 6.51 |
| KIF53_RS20415 | acyltransferase domain-containing protein | 6.44 |
| KIF53_RS06810 | cyanide-forming glycine dehydrogenase subunit HcnA | 6.33 |
| KIF53_RS21020 | PfaD family polyunsaturated fatty acid/polyketide biosynthesis protein | 6.18 |
| KIF53_RS14555 | DUF1842 domain-containing protein | 6.18 |
| KIF53_RS01860 | long-chain-fatty-acyl-CoA reductase | 5.94 |
| KIF53_RS21015 | DUF3149 domain-containing protein | 5.88 |
| KIF53_RS19385 | amino acid adenylation domain-containing protein | 5.87 |
| KIF53_RS15570 | hypothetical protein | 5.74 |
| KIF53_RS01870 | MdfA family multidrug efflux MFS transporter | 5.53 |
| KIF53_RS14680 | U32 family peptidase | 5.47 |
| KIF53_RS06800 | cyanide-forming glycine dehydrogenase subunit HcnC | 5.44 |
| KIF53_RS05635 | FAD-dependent oxidoreductase | 5.42 |
| KIF53_RS16820 | M4 family metallopeptidase | 5.35 |
| KIF53_RS16340 | ornithine carbamoyltransferase | 5.23 |
| KIF53_RS19375 | ACP S-malonyltransferase | 5.23 |
| KIF53_RS05645 | FAD-dependent monooxygenase | 5.21 |
| KIF53_RS11595 | bifunctional phosphoribosylaminoimidazolecarboxamide formyltransferase/IMP cyclohydrolase | 5.08 |
| KIF53_RS05655 | violacein biosynthesis enzyme VioE | 5.04 |
| KIF53_RS13295 | hypothetical protein | 4.99 |
| KIF53_RS15685 | protein | 4.97 |
| KIF53_RS14675 | SCP2 sterol-binding domain-containing protein | 4.94 |
| KIF53_RS19390 | acyl-CoA dehydrogenase family protein | 4.87 |
| KIF53_RS21040 | hypothetical protein | 4.86 |
| KIF53_RS16335 | arginine deiminase | 4.86 |
| KIF53_RS21035 | polysaccharide pyruvyl transferase family protein | 4.82 |
| KIF53_RS01145 | hypothetical protein | 4.81 |
| KIF53_RS21030 | 4'-phosphopantetheinyl transferase superfamily protein | 4.74 |
| KIF53_RS21025 | alpha/beta fold hydrolase | 4.74 |
| KIF53_RS19370 | AMP-binding protein | 4.73 |
| KIF53_RS06795 | ABC transporter substrate-binding protein | 4.71 |

| | | |
|---|---|---|
| KIF53_RS05640 | iminophenyl-pyruvate dimer synthase VioB | 4.71 |
| KIF53_RS19365 | type I polyketide synthase | 4.53 |
| KIF53_RS05650 | tryptophan hydroxylase | 4.52 |
| KIF53_RS01125 | hypothetical protein | 4.52 |
| KIF53_RS01855 | AMP-dependent synthetase | 4.50 |
| KIF53_RS02825 | hydrogenase maturation protein | 4.48 |
| KIF53_RS15785 | methyl-accepting chemotaxis protein | 4.45 |
| KIF53_RS14685 | U32 family peptidase | 4.42 |
| KIF53_RS17975 | aspartate carbamoyltransferase | 4.34 |
| KIF53_RS16345 | carbamate kinase | 4.33 |
| KIF53_RS16060 | adenylosuccinate lyase | 4.32 |
| KIF53_RS06780 | ABC transporter ATP-binding protein | 4.31 |
| KIF53_RS16225 | carbamoyl-phosphate synthase large subunit | 4.29 |
| KIF53_RS13290 | hypothetical protein | 4.26 |
| KIF53_RS06785 | ABC transporter permease subunit | 4.25 |
| KIF53_RS19360 | serine hydrolase | 4.23 |
| KIF53_RS08980 | acetyl-CoA carboxylase biotin carboxyl carrier protein | 4.16 |
| KIF53_RS10155 | amidophosphoribosyltransferase | 4.12 |
| KIF53_RS19380 | AMP-binding protein | 4.08 |
| KIF53_RS19395 | acyl carrier protein | 4.07 |
| KIF53_RS01120 | glycosyltransferase family 2 protein | 4.02 |
| KIF53_RS03970 | type I-E CRISPR-associated protein Cas6/Cse3/CasE | 4.02 |
| KIF53_RS13300 | peptidase | 4.00 |
| KIF53_RS05185 | sulfur subunit | 3.99 |
| KIF53_RS16220 | leucine efflux protein LeuE | 3.91 |
| KIF53_RS13815 | helix-turn-helix domain-containing protein | 3.90 |
| KIF53_RS21000 | hypothetical protein | 3.82 |
| KIF53_RS11705 | helix-turn-helix domain-containing protein | 3.74 |
| KIF53_RS05740 | hypothetical protein | 3.72 |
| KIF53_RS05695 | cold-shock protein | 3.71 |
| KIF53_RS17075 | porin | 3.70 |
| KIF53_RS15420 | HNH endonuclease | 3.64 |
| KIF53_RS15405 | hypothetical protein | 3.58 |
| KIF53_RS03965 | type I-E CRISPR-associated protein Cas5/CasD | 3.56 |
| KIF53_RS01130 | transketolase | 3.54 |
| KIF53_RS00815 | type I glyceraldehyde-3-phosphate dehydrogenase | 3.52 |
| KIF53_RS18720 | dehydrogenase/methenyltetrahydrofolate cyclohydrolase FolD | 3.52 |
| KIF53_RS19350 | AMP-binding protein | 3.50 |
| KIF53_RS13245 | PAAR domain-containing protein | 3.47 |
| KIF53_RS13360 | M15 family metallopeptidase | 3.46 |
| KIF53_RS07505 | flagellar motor switch protein FliN | 3.45 |
| KIF53_RS03945 | CRISPR-associated helicase/endonuclease Cas3 | 3.45 |
| KIF53_RS04005 | L-serine ammonia-lyase | 3.45 |
| KIF53_RS09805 | hypothetical protein | 3.44 |

| | | |
|---|---|---|
| KIF53_RS11695 | chorismate-binding protein | 3.43 |
| KIF53_RS19355 | amino acid adenylation domain-containing protein | 3.43 |
| KIF53_RS06790 | ABC transporter permease subunit | 3.42 |
| KIF53_RS03950 | type I-E CRISPR-associated protein Cse1/CasA | 3.41 |
| KIF53_RS13260 | phage major tail tube protein | 3.38 |
| KIF53_RS19335 | alpha/beta fold hydrolase | 3.36 |
| KIF53_RS00650 | 5-(carboxyamino)imidazole ribonucleotide mutase | 3.34 |
| KIF53_RS20220 | chaperonin GroEL | 3.34 |
| KIF53_RS20765 | indolepyruvate ferredoxin oxidoreductase family protein | 3.32 |
| KIF53_RS03960 | type I-E CRISPR-associated protein Cas7/Cse4/CasC | 3.32 |
| KIF53_RS00655 | 5-(carboxyamino)imidazole ribonucleotide synthase | 3.30 |
| KIF53_RS15110 | P-II family nitrogen regulator | 3.30 |
| KIF53_RS02575 | elongation factor G | 3.28 |
| KIF53_RS18405 | MarR family transcriptional regulator | 3.28 |
| KIF53_RS04885 | aminomethyl-transferring glycine dehydrogenase | 3.28 |
| KIF53_RS07155 | oxidoreductase | 3.28 |
| KIF53_RS01150 | subunit | 3.27 |
| KIF53_RS16385 | NCS2 family permease | 3.23 |
| KIF53_RS04965 | acyl carrier protein | 3.20 |
| KIF53_RS17070 | FAD-dependent oxidoreductase | 3.20 |
| KIF53_RS07315 | integration host factor subunit beta | 3.19 |
| KIF53_RS10620 | DUF3313 family protein | 3.19 |
| KIF53_RS02970 | hypothetical protein | 3.19 |
| KIF53_RS14205 | iron transporter | 3.18 |
| KIF53_RS01535 | septal ring lytic transglycosylase RlpA family protein | 3.16 |
| KIF53_RS07500 | flagellar type III secretion system pore protein FliP | 3.15 |
| KIF53_RS12190 | F0F1 ATP synthase subunit beta | 3.14 |
| KIF53_RS00670 | synthase | 3.14 |
| KIF53_RS13265 | phage tail sheath subtilisin-like domain-containing protein | 3.12 |
| KIF53_RS21590 | DUF6531 domain-containing protein | 3.11 |
| KIF53_RS21840 | DUF805 domain-containing protein | 3.10 |
| KIF53_RS18410 | energy transducer TonB | 3.10 |
| KIF53_RS01115 | glycosyltransferase | 3.09 |
| KIF53_RS12180 | F0F1 ATP synthase subunit alpha | 3.09 |
| KIF53_RS15065 | co-chaperone GroES | 3.08 |
| KIF53_RS19340 | cupin-like domain-containing protein | 3.08 |
| KIF53_RS02670 | 50S ribosomal protein L18 | 3.08 |
| KIF53_RS15070 | chaperonin GroEL | 3.06 |
| KIF53_RS21585 | DUF2345 domain-containing protein | 3.06 |
| KIF53_RS13270 | hypothetical protein | 3.06 |
| KIF53_RS02580 | elongation factor Tu | 3.03 |
| KIF53_RS20225 | co-chaperone GroES | 3.00 |
| KIF53_RS04740 | phosphopyruvate hydratase | 2.99 |
| KIF53_RS12185 | F0F1 ATP synthase subunit gamma | 2.98 |
| KIF53_RS17825 | energy transducer TonB | 2.97 |

| | | |
|---|---|---|
| KIF53_RS01140 | hypothetical protein | 2.96 |
| KIF53_RS07310 | 30S ribosomal protein S1 | 2.96 |
| KIF53_RS05660 | hypothetical protein | 2.95 |
| KIF53_RS08985 | acetyl-CoA carboxylase biotin carboxylase subunit | 2.94 |
| KIF53_RS13285 | DUF2190 family protein | 2.94 |
| KIF53_RS02680 | 50S ribosomal protein L30 | 2.94 |
| KIF53_RS02675 | 30S ribosomal protein S5 | 2.93 |
| KIF53_RS16215 | small subunit | 2.93 |
| KIF53_RS17940 | DUF2059 domain-containing protein | 2.93 |
| KIF53_RS05180 | fumarate reductase (quinol) flavoprotein subunit | 2.93 |
| KIF53_RS20390 | methyltransferase domain-containing protein | 2.91 |
| KIF53_RS14110 | GspH/FimT family pseudopilin | 2.90 |
| KIF53_RS02540 | tRNA-Thr | 2.90 |
| KIF53_RS12560 | BMP family ABC transporter substrate-binding protein | 2.90 |
| KIF53_RS03535 | type 1 fimbrial protein | 2.89 |
| KIF53_RS02610 | 30S ribosomal protein S19 | 2.89 |
| KIF53_RS02665 | 50S ribosomal protein L6 | 2.89 |
| KIF53_RS01110 | acyltransferase | 2.88 |
| KIF53_RS02830 | acyl-CoA dehydrogenase family protein | 2.87 |
| KIF53_RS12175 | F0F1 ATP synthase subunit delta | 2.87 |
| KIF53_RS17840 | biopolymer transporter ExbD | 2.87 |
| KIF53_RS07495 | flagellar biosynthesis protein FliQ | 2.86 |
| KIF53_RS05190 | fumarate reductase | 2.84 |
| KIF53_RS17680 | NERD domain-containing protein | 2.84 |
| KIF53_RS13280 | DUF1320 domain-containing protein | 2.82 |
| KIF53_RS04880 | glycine cleavage system protein GcvH | 2.82 |
| KIF53_RS15870 | phosphoribosylformylglycinamidine synthase | 2.81 |
| KIF53_RS11700 | queuosine precursor transporter | 2.81 |
| KIF53_RS04205 | cytochrome c oxidase accessory protein CcoG | 2.80 |
| KIF53_RS08500 | hypothetical protein | 2.80 |
| KIF53_RS05055 | cystathionine beta-synthase | 2.78 |
| KIF53_RS01540 | tRNA (uridine(34)/cytosine(34)/5-carboxymethylaminomethyluridine(34)-2'-O)-methyltransferase TrmL | 2.78 |
| KIF53_RS06905 | flagellar hook-basal body complex protein | 2.78 |
| KIF53_RS02660 | 30S ribosomal protein S8 | 2.78 |
| KIF53_RS05060 | protein | 2.77 |
| KIF53_RS02545 | 50S ribosomal protein L10 | 2.74 |
| KIF53_RS02270 | orotate phosphoribosyltransferase | 2.73 |
| KIF53_RS22035 | TerD family protein | 2.71 |
| KIF53_RS13250 | hypothetical protein | 2.70 |
| KIF53_RS19325 | 3-hydroxyacyl-CoA dehydrogenase family protein | 2.70 |
| KIF53_RS13330 | DUF3486 family protein | 2.70 |
| KIF53_RS02595 | 50S ribosomal protein L4 | 2.70 |
| KIF53_RS11445 | hypothetical protein | 2.70 |

| | | |
|---|---|---|
| KIF53_RS00660 | SEL1-like repeat protein | 2.70 |
| KIF53_RS21790 | hypothetical protein | 2.70 |
| KIF53_RS12165 | F0F1 ATP synthase subunit C | 2.68 |
| KIF53_RS02975 | glycosyltransferase family 4 protein | 2.67 |
| KIF53_RS02605 | 50S ribosomal protein L2 | 2.67 |
| KIF53_RS16955 | fucose-binding lectin | 2.66 |
| KIF53_RS12170 | F0F1 ATP synthase subunit B | 2.66 |
| KIF53_RS02655 | 30S ribosomal protein S14 | 2.65 |
| KIF53_RS05525 | carbohydrate porin | 2.65 |
| KIF53_RS09800 | activating protein | 2.64 |
| KIF53_RS01980 | hypothetical protein | 2.64 |
| KIF53_RS08190 | nuclear transport factor 2 family protein | 2.63 |
| KIF53_RS04970 | beta-ketoacyl-ACP synthase II | 2.63 |
| KIF53_RS14595 | rhodoquinone biosynthesis methyltransferase RquA | 2.62 |
| KIF53_RS01155 | peptidase domain-containing ABC transporter | 2.61 |
| KIF53_RS07480 | flagellar biosynthesis protein FlhA | 2.61 |
| KIF53_RS06910 | flagellar basal body rod protein FlgF | 2.60 |
| KIF53_RS08490 | ADP-forming succinate--CoA ligase subunit beta | 2.59 |
| KIF53_RS05030 | (2Fe-2S)-binding protein | 2.59 |
| KIF53_RS06960 | cytochrome c | 2.59 |
| KIF53_RS06000 | tRNA-Asp | 2.58 |
| KIF53_RS00960 | proton-translocating transhydrogenase family protein | 2.58 |
| KIF53_RS11805 | DUF2860 family protein | 2.58 |
| KIF53_RS04950 | ketoacyl-ACP synthase III | 2.57 |
| KIF53_RS02615 | 50S ribosomal protein L22 | 2.56 |
| KIF53_RS09795 | anaerobic ribonucleoside-triphosphate reductase | 2.55 |
| KIF53_RS06895 | flagellar basal body rod protein FlgC | 2.55 |
| KIF53_RS11255 | glutamate--ammonia ligase | 2.55 |
| KIF53_RS06310 | GNAT family N-acetyltransferase | 2.54 |
| KIF53_RS03975 | type I-E CRISPR-associated endonuclease Cas1e | 2.53 |
| KIF53_RS21355 | Smr/MutS family protein | 2.53 |
| KIF53_RS03060 | outer membrane beta-barrel protein | 2.53 |
| KIF53_RS21595 | hypothetical protein | 2.53 |
| KIF53_RS02880 | class I SAM-dependent methyltransferase | 2.52 |
| KIF53_RS17440 | UDP-2,3-diacylglucosamine diphosphatase | 2.50 |
| KIF53_RS02035 | acid phosphatase | 2.47 |
| KIF53_RS02550 | 50S ribosomal protein L7/L12 | 2.46 |
| KIF53_RS10935 | GbsR/MarR family transcriptional regulator | 2.46 |
| KIF53_RS06900 | flagellar basal body rod modification protein FlgD | 2.46 |
| KIF53_RS06055 | cobyrinate a,c-diamide synthase | 2.46 |
| KIF53_RS16910 | GyrI-like domain-containing protein | 2.45 |
| KIF53_RS08485 | succinate--CoA ligase subunit alpha | 2.45 |
| KIF53_RS04195 | LysR family transcriptional regulator | 2.44 |
| KIF53_RS10945 | cytochrome d ubiquinol oxidase subunit II | 2.44 |
| KIF53_RS21290 | adenylosuccinate synthase | 2.44 |

| | | |
|---|---|---|
| KIF53_RS13745 | acetyl-CoA C-acyltransferase | 2.43 |
| KIF53_RS02885 | sulfotransferase | 2.43 |
| KIF53_RS12915 | BMP family ABC transporter substrate-binding protein | 2.42 |
| KIF53_RS04010 | HAAAP family serine/threonine permease | 2.42 |
| KIF53_RS03980 | type I-E CRISPR-associated endoribonuclease Cas2 | 2.41 |
| KIF53_RS06660 | protoporphyrinogen oxidase | 2.41 |
| KIF53_RS13740 | MerR family DNA-binding transcriptional regulator | 2.41 |
| KIF53_RS00665 | DNA alkylation repair protein | 2.40 |
| KIF53_RS19725 | enterobactin transporter EntS | 2.40 |
| KIF53_RS15105 | ammonium transporter | 2.39 |
| KIF53_RS19960 | nicotinate-nucleotide adenylyltransferase | 2.39 |
| KIF53_RS18715 | tRNA-Pro | 2.39 |
| KIF53_RS08975 | type II 3-dehydroquinate dehydratase | 2.38 |
| KIF53_RS05195 | fumarate reductase subunit FrdD | 2.38 |
| KIF53_RS02535 | 50S ribosomal protein L1 | 2.36 |
| KIF53_RS05475 | 2OG-Fe dioxygenase family protein | 2.36 |
| KIF53_RS11710 | hypothetical protein | 2.36 |
| KIF53_RS02600 | 50S ribosomal protein L23 | 2.36 |
| KIF53_RS10510 | QueF | 2.36 |
| KIF53_RS04285 | polyribonucleotide nucleotidyltransferase | 2.35 |
| KIF53_RS12815 | hypothetical protein | 2.34 |
| KIF53_RS16905 | tetratricopeptide repeat protein | 2.33 |
| KIF53_RS02935 | DUF4337 domain-containing protein | 2.33 |
| KIF53_RS07055 | hypothetical protein | 2.33 |
| KIF53_RS02530 | 50S ribosomal protein L11 | 2.33 |
| KIF53_RS09770 | NAD(P)H-quinone oxidoreductase | 2.32 |
| KIF53_RS04875 | glycine cleavage system aminomethyltransferase GcvT | 2.32 |
| KIF53_RS15835 | DUF697 domain-containing protein | 2.32 |
| KIF53_RS04340 | iron ABC transporter permease | 2.31 |
| KIF53_RS02690 | preprotein translocase subunit SecY | 2.30 |
| KIF53_RS03905 | EAL domain-containing protein | 2.29 |
| KIF53_RS06925 | flagellar basal body P-ring protein FlgI | 2.29 |
| KIF53_RS10940 | cytochrome ubiquinol oxidase subunit I | 2.27 |
| KIF53_RS02715 | 30S ribosomal protein S4 | 2.27 |
| KIF53_RS07530 | flagellar assembly protein H | 2.26 |
| KIF53_RS17425 | OmpP1/FadL family transporter | 2.26 |
| KIF53_RS11770 | DUF3619 family protein | 2.25 |
| KIF53_RS09615 | hypothetical protein | 2.25 |
| KIF53_RS02950 | oligosaccharide flippase family protein | 2.25 |
| KIF53_RS07535 | flagellar protein export ATPase FliI | 2.25 |
| KIF53_RS11060 | anaerobic C4-dicarboxylate transporter | 2.24 |
| KIF53_RS02620 | 30S ribosomal protein S3 | 2.24 |
| KIF53_RS07515 | flagellar hook-basal body complex protein FliE | 2.24 |
| KIF53_RS12195 | F0F1 ATP synthase subunit epsilon | 2.24 |
| KIF53_RS04945 | phosphate acyltransferase PlsX | 2.23 |

| | | |
|---|---|---|
| KIF53_RS17835 | MotA/TolQ/ExbB proton channel family protein | 2.23 |
| KIF53_RS13735 | 3-hydroxybutyryl-CoA dehydrogenase | 2.22 |
| KIF53_RS02960 | glycosyltransferase family 4 protein | 2.22 |
| KIF53_RS17140 | hypothetical protein | 2.22 |
| KIF53_RS08495 | dihydrolipoyl dehydrogenase | 2.22 |
| KIF53_RS19555 | phosphonate C-P lyase system protein PhnG | 2.22 |
| KIF53_RS02310 | chitinase | 2.21 |
| KIF53_RS02870 | sulfotransferase family 2 domain-containing protein | 2.21 |
| KIF53_RS04350 | DUF2218 domain-containing protein | 2.21 |
| KIF53_RS06915 | flagellar basal-body rod protein FlgG | 2.20 |
| KIF53_RS05610 | hypothetical protein | 2.20 |
| KIF53_RS02570 | 30S ribosomal protein S7 | 2.20 |
| KIF53_RS07120 | NAD-glutamate dehydrogenase | 2.19 |
| KIF53_RS09855 | phosphatidate cytidylyltransferase | 2.19 |
| KIF53_RS01850 | GNAT family N-acetyltransferase | 2.19 |
| KIF53_RS01635 | copper chaperone PCu(A)C | 2.19 |
| KIF53_RS10515 | CPBP family intramembrane metalloprotease | 2.19 |
| KIF53_RS06890 | flagellar basal body rod protein FlgB | 2.18 |
| KIF53_RS16005 | hypothetical protein | 2.17 |
| KIF53_RS22550 | type VI secretion system tip protein VgrG | 2.17 |
| KIF53_RS02500 | tRNA-Gly | 2.17 |
| KIF53_RS02945 | DegT/DnrJ/EryC1/StrS family aminotransferase | 2.17 |
| KIF53_RS15775 | metal-dependent hydrolase | 2.16 |
| KIF53_RS05745 | sugar MFS transporter | 2.15 |
| KIF53_RS15825 | Ldh family oxidoreductase | 2.15 |
| KIF53_RS09860 | 1-deoxy-D-xylulose-5-phosphate reductoisomerase | 2.15 |
| KIF53_RS02720 | DNA-directed RNA polymerase subunit alpha | 2.14 |
| KIF53_RS09905 | DNA-deoxyinosine glycosylase | 2.14 |
| KIF53_RS22040 | VWA domain-containing protein | 2.14 |
| KIF53_RS06060 | cob(I)yrinic acid a,c-diamide adenosyltransferase | 2.13 |
| KIF53_RS07555 | hypothetical protein | 2.13 |
| KIF53_RS00575 | isopenicillin N synthase family oxygenase | 2.13 |
| KIF53_RS07440 | ribose ABC transporter substrate-binding protein RbsB | 2.13 |
| KIF53_RS11335 | DUF934 domain-containing protein | 2.12 |
| KIF53_RS13275 | Gp37 family protein | 2.12 |
| KIF53_RS02965 | glycosyltransferase family 4 protein | 2.12 |
| KIF53_RS06085 | precorrin-3B C(17)-methyltransferase | 2.11 |
| KIF53_RS06095 | precorrin-4 C(11)-methyltransferase | 2.11 |
| KIF53_RS00965 | beta | 2.11 |
| KIF53_RS13645 | metal-dependent hydrolase | 2.11 |
| KIF53_RS12160 | F0F1 ATP synthase subunit A | 2.10 |
| KIF53_RS01355 | hypothetical protein | 2.10 |
| KIF53_RS07540 | flagellar export protein FliJ | 2.10 |
| KIF53_RS02890 | tRNA-Met | 2.10 |
| KIF53_RS12920 | MATE family efflux transporter | 2.10 |

| | | |
|---|---|---|
| KIF53_RS15815 | hypothetical protein | 2.09 |
| KIF53_RS11435 | dihydroorotate oxidase | 2.09 |
| KIF53_RS12570 | ABC transporter permease | 2.09 |
| KIF53_RS20600 | acyl-CoA thioesterase | 2.09 |
| KIF53_RS22580 | type VI secretion system Vgr family protein | 2.09 |
| KIF53_RS22005 | hypothetical protein | 2.09 |
| KIF53_RS12575 | ABC transporter permease | 2.08 |
| KIF53_RS13730 | dehydrogenase | 2.08 |
| KIF53_RS04335 | ABC transporter ATP-binding protein | 2.08 |
| KIF53_RS00820 | transketolase | 2.08 |
| KIF53_RS10395 | peptidoglycan DD-metalloendopeptidase family protein | 2.07 |
| KIF53_RS02505 | tRNA-Thr | 2.07 |
| KIF53_RS07685 | sigma-70 family RNA polymerase sigma factor | 2.07 |
| KIF53_RS06065 | iron ABC transporter permease | 2.06 |
| KIF53_RS05240 | DUF456 family protein | 2.06 |
| KIF53_RS04960 | 3-oxoacyl-ACP reductase FabG | 2.06 |
| KIF53_RS07485 | flagellar type III secretion system protein FlhB | 2.05 |
| KIF53_RS22030 | hypothetical protein | 2.05 |
| KIF53_RS16885 | nuclear transport factor 2 family protein | 2.05 |
| KIF53_RS16330 | arginine-ornithine antiporter | 2.05 |
| KIF53_RS20805 | glutathione transferase GstA | 2.05 |
| KIF53_RS03955 | type I-E CRISPR-associated protein Cse2/CasB | 2.04 |
| KIF53_RS11450 | catalase | 2.04 |
| KIF53_RS03920 | elongation factor P | 2.04 |
| KIF53_RS16605 | phosphatase PAP2 family protein | 2.04 |
| KIF53_RS16020 | formate dehydrogenase accessory protein FdhE | 2.04 |
| KIF53_RS07520 | flagellar M-ring protein FliF | 2.03 |
| KIF53_RS19230 | MFS transporter | 2.03 |
| KIF53_RS13650 | holo-ACP synthase | 2.03 |
| KIF53_RS02650 | 50S ribosomal protein L5 | 2.03 |
| KIF53_RS09760 | cbb3-type cytochrome oxidase assembly protein CcoS | 2.03 |
| KIF53_RS04355 | TonB-dependent receptor | 2.03 |
| KIF53_RS06580 | nucleotide exchange factor GrpE | 2.03 |
| KIF53_RS21785 | hypothetical protein | 2.02 |
| KIF53_RS05680 | aminotransferase class III-fold pyridoxal phosphate-dependent enzyme | 2.02 |
| KIF53_RS06080 | GTP-binding protein | 2.02 |
| KIF53_RS07525 | flagellar motor switch protein FliG | 2.02 |
| KIF53_RS08250 | methyl-accepting chemotaxis protein | 2.02 |
| KIF53_RS16130 | MaoC family dehydratase | 2.01 |
| KIF53_RS03760 | preprotein translocase subunit YajC | 2.01 |
| KIF53_RS01520 | cytochrome c biogenesis protein ResB | 2.01 |
| KIF53_RS02980 | DegT/DnrJ/EryC1/StrS family aminotransferase | 2.01 |
| KIF53_RS02925 | YdcF family protein | 2.00 |

### *C. subtsugae* genes activated by C8-HSL

| Locus tag corresponding with accession NZ_JAHDTB000000000.1 | Predicted gene function | $Log_2$ fold change compared with no AHLs |
|---|---|---|
| KIF53_RS16340 | cyanide-forming glycine dehydrogenase subunit HcnA | 6.41 |
| KIF53_RS20400 | cyanide-forming glycine dehydrogenase subunit HcnC | 5.61 |
| KIF53_RS02310 | ornithine carbamoyltransferase | 5.22 |
| KIF53_RS06310 | hybrid non-ribosomal peptide synthetase/type I polyketide synthase | 4.97 |
| KIF53_RS15105 | chitinase | 4.71 |
| KIF53_RS06795 | GNAT family N-acetyltransferase | 3.86 |
| KIF53_RS20405 | ammonium transporter | 3.97 |
| KIF53_RS21025 | ABC transporter substrate-binding protein | 4.58 |
| KIF53_RS06805 | amino acid adenylation domain-containing protein | 4.80 |
| KIF53_RS05635 | alpha/beta fold hydrolase | 4.65 |
| KIF53_RS15685 | NAD(P)/FAD-dependent oxidoreductase | 6.95 |
| KIF53_RS05525 | FAD-dependent oxidoreductase | 4.14 |
| KIF53_RS16345 | M9 family metallopeptidase N-terminal domain-containing protein | 3.67 |
| KIF53_RS04205 | carbohydrate porin | 3.16 |
| KIF53_RS14555 | carbamate kinase | 3.93 |
| KIF53_RS13810 | cytochrome c oxidase accessory protein CcoG | 3.35 |
| KIF53_RS16335 | hybrid non-ribosomal peptide synthetase/type I polyketide synthase | 3.29 |
| KIF53_RS06780 | activator protein | 4.40 |
| KIF53_RS02825 | arginine deiminase | 3.95 |
| KIF53_RS20410 | ABC transporter ATP-binding protein | 4.43 |
| KIF53_RS21040 | hydrogenase maturation protein | 3.18 |
| KIF53_RS21035 | non-ribosomal peptide synthetase | 4.04 |
| KIF53_RS15785 | hypothetical protein | 4.75 |
| KIF53_RS15110 | polysaccharide pyruvyl transferase family protein | 3.91 |
| KIF53_RS05190 | methyl-accepting chemotaxis protein | 2.60 |
| KIF53_RS05180 | P-II family nitrogen regulator | 3.83 |
| KIF53_RS11805 | fumarate reductase | 3.17 |
| KIF53_RS17975 | fumarate reductase (quinol) flavoprotein subunit | 2.70 |
| KIF53_RS14595 | DUF2860 family protein | 2.53 |
| KIF53_RS01130 | aspartate carbamoyltransferase | 3.03 |
| KIF53_RS03060 | rhodoquinone biosynthesis methyltransferase RquA | 2.70 |
| KIF53_RS16905 | transketolase | 3.08 |
| KIF53_RS21030 | outer membrane beta-barrel protein | 3.38 |
| KIF53_RS10620 | tetratricopeptide repeat protein | 2.60 |

| | | |
|---|---|---|
| KIF53_RS18600 | 4'-phosphopantetheinyl transferase superfamily protein | 3.58 |
| KIF53_RS10945 | DUF3313 family protein | 2.82 |
| KIF53_RS01150 | LysE family translocator | 2.91 |
| KIF53_RS13710 | cytochrome d ubiquinol oxidase subunit II | 3.05 |
| KIF53_RS17525 | HlyD family efflux transporter periplasmic adaptor subunit | 2.95 |
| KIF53_RS13945 | 3-hydroxyisobutyrate dehydrogenase | 2.29 |
| KIF53_RS13960 | ssDNA-binding domain-containing protein | 3.06 |
| KIF53_RS13880 | hypothetical protein | 2.65 |
| KIF53_RS16225 | Flp pilus assembly complex ATPase component TadA | 2.41 |
| KIF53_RS13925 | hypothetical protein | 2.61 |
| KIF53_RS05195 | carbamoyl-phosphate synthase large subunit | 2.71 |
| KIF53_RS17485 | PilN family type IVB pilus formation outer membrane protein | 2.66 |
| KIF53_RS17680 | fumarate reductase subunit FrdD | 2.88 |
| KIF53_RS09615 | DotA/TraY family protein | 2.45 |
| KIF53_RS20765 | NERD domain-containing protein | 2.66 |
| KIF53_RS11255 | hypothetical protein | 3.10 |
| KIF53_RS20415 | indolepyruvate ferredoxin oxidoreductase family protein | 2.52 |
| KIF53_RS20465 | glutamate--ammonia ligase | 3.21 |
| KIF53_RS10940 | acyltransferase domain-containing protein | 3.73 |
| KIF53_RS01145 | GNAT family N-acetyltransferase | 2.46 |
| KIF53_RS17060 | cytochrome ubiquinol oxidase subunit I | 2.82 |
| KIF53_RS16910 | hypothetical protein | 3.93 |
| KIF53_RS17075 | magnesium-translocating P-type ATPase | 2.47 |
| KIF53_RS05185 | GyrI-like domain-containing protein | 2.86 |
| KIF53_RS11400 | porin | 3.25 |
| KIF53_RS13855 | succinate dehydrogenase/fumarate reductase iron-sulfur subunit | 4.07 |
| KIF53_RS00720 | histone deacetylase family protein | 2.05 |
| KIF53_RS21000 | hypothetical protein | 2.44 |
| KIF53_RS01125 | depolymerase | 2.77 |
| KIF53_RS18595 | hypothetical protein | 2.96 |
| KIF53_RS14025 | hypothetical protein | 3.15 |
| KIF53_RS21015 | PLP-dependent aminotransferase family protein | 2.36 |
| KIF53_RS15070 | YeiH family putative sulfate export transporter | 2.21 |
| KIF53_RS07545 | hypothetical protein | 3.13 |
| KIF53_RS12920 | chaperonin GroEL | 2.84 |
| KIF53_RS13860 | flagellar filament capping protein FliD | 2.22 |
| KIF53_RS20390 | MATE family efflux transporter | 2.30 |
| KIF53_RS17750 | hypothetical protein | 2.47 |
| KIF53_RS21840 | methyltransferase domain-containing protein | 2.79 |
| KIF53_RS04885 | L-threonine dehydrogenase | 2.50 |
| KIF53_RS17520 | DUF805 domain-containing protein | 2.61 |
| KIF53_RS05640 | aminomethyl-transferring glycine dehydrogenase | 2.40 |

| | | |
|---|---|---|
| KIF53_RS06790 | hypothetical protein | 3.22 |
| KIF53_RS13715 | iminophenyl-pyruvate dimer synthase VioB | 2.51 |
| KIF53_RS15405 | ABC transporter permease subunit | 2.81 |
| KIF53_RS16915 | enoyl-CoA hydratase/isomerase family protein | 2.29 |
| KIF53_RS19365 | hypothetical protein | 2.50 |
| KIF53_RS16820 | hydroxymethylglutaryl-CoA lyase | 2.24 |
| KIF53_RS13920 | type I polyketide synthase | 2.56 |
| KIF53_RS21010 | M4 family metallopeptidase | 2.76 |
| KIF53_RS01140 | toxin co-regulated pilus biosynthesis Q family protein | 2.87 |
| KIF53_RS01120 | SDR family NAD(P)-dependent oxidoreductase | 4.10 |
| KIF53_RS00965 | hypothetical protein | 2.56 |
| KIF53_RS04020 | glycosyltransferase family 2 protein | 2.89 |
| KIF53_RS17515 | NAD(P)(+) transhydrogenase (Re/Si-specific) subunit beta | 2.03 |
| KIF53_RS07155 | formate C-acetyltransferase | 2.20 |
| KIF53_RS01860 | polymer-forming cytoskeletal protein | 2.34 |
| KIF53_RS06785 | oxidoreductase | 2.73 |
| KIF53_RS09620 | long-chain-fatty-acyl-CoA reductase | 2.10 |
| KIF53_RS16955 | ABC transporter permease subunit | 3.49 |
| KIF53_RS16900 | beta-ketoacyl-ACP synthase | 2.24 |
| KIF53_RS19375 | fucose-binding lectin | 2.10 |
| KIF53_RS13940 | acetyl/propionyl/methylcrotonyl-CoA carboxylase subunit alpha | 2.07 |
| KIF53_RS00955 | ACP S-malonyltransferase | 2.69 |
| KIF53_RS01960 | Flp pilus assembly complex ATPase component TadA | 2.51 |
| KIF53_RS17530 | Re/Si-specific NAD(P)(+) transhydrogenase subunit alpha | 2.08 |
| KIF53_RS15570 | universal stress protein | 2.36 |
| KIF53_RS01110 | hypothetical protein | 2.44 |
| KIF53_RS03105 | hypothetical protein | 2.35 |
| KIF53_RS19370 | acyltransferase | 2.40 |
| KIF53_RS12190 | cbb3-type cytochrome c oxidase subunit 3 | 2.12 |
| KIF53_RS13895 | AMP-binding protein | 2.55 |
| KIF53_RS13900 | F0F1 ATP synthase subunit beta | 2.39 |
| KIF53_RS13930 | type IV secretory system conjugative DNA transfer family protein | 2.42 |
| KIF53_RS16325 | Flp pilus assembly complex ATPase component TadA | 2.47 |
| KIF53_RS14685 | type 4b pilus protein PilO2 | 2.61 |
| KIF53_RS13955 | DUF3149 domain-containing protein | 3.74 |
| KIF53_RS09025 | U32 family peptidase | 2.25 |
| KIF53_RS13815 | hypothetical protein | 2.09 |
| KIF53_RS08485 | PTS glucose transporter subunit IIBC | 2.07 |
| KIF53_RS09020 | helix-turn-helix domain-containing protein | 2.01 |
| KIF53_RS05540 | succinate--CoA ligase subunit alpha | 2.13 |
| KIF53_RS02580 | phosphoenolpyruvate--protein phosphotransferase | 2.06 |

| | | |
|---|---|---|
| KIF53_RS00960 | hypothetical protein | 2.10 |
| KIF53_RS01115 | elongation factor Tu | 2.23 |
| KIF53_RS21020 | proton-translocating transhydrogenase family protein | 2.39 |
| KIF53_RS13865 | glycosyltransferase | 2.04 |
| KIF53_RS15065 | PfaD family polyunsaturated fatty acid/polyketide biosynthesis protein | 2.97 |
| KIF53_RS13730 | hypothetical protein | 3.05 |
| KIF53_RS07290 | co-chaperone GroES | 2.24 |
| KIF53_RS22565 | CoA-acylating methylmalonate-semialdehyde dehydrogenase | 2.07 |
| KIF53_RS15830 | cytochrome b | 2.10 |
| KIF53_RS22325 | hypothetical protein | 2.28 |
| KIF53_RS17610 | hypothetical protein | 2.34 |
| KIF53_RS06960 | hypothetical protein | 2.39 |
| KIF53_RS05655 | phenylacetate-CoA oxygenase subunit PaaI | 2.15 |
| KIF53_RS09625 | cytochrome c | 2.05 |
| KIF53_RS03110 | violacein biosynthesis enzyme VioE | 2.33 |
| KIF53_RS13875 | 3-oxoacyl-ACP reductase FabG | 2.80 |
| KIF53_RS19360 | cytochrome-c oxidase, cbb3-type subunit II | 2.03 |
| KIF53_RS21790 | hypothetical protein | 2.42 |
| KIF53_RS13850 | serine hydrolase | 2.09 |
| KIF53_RS22560 | hypothetical protein | 2.12 |
| KIF53_RS09630 | DotI/IcmL family type IV secretion protein | 2.22 |
| KIF53_RS19555 | hypothetical protein | 2.08 |
| KIF53_RS14680 | hotdog family protein | 2.43 |
| KIF53_RS12980 | phosphonate C-P lyase system protein PhnG | 2.01 |
| KIF53_RS05645 | U32 family peptidase | 2.32 |
| KIF53_RS02970 | type III secretion system inner rod subunit SctI | 2.01 |
| KIF53_RS14110 | FAD-dependent monooxygenase | 2.22 |
| | hypothetical protein | 2.07 |
| | GspH/FimT family pseudopilin | 2.02 |
